# Decoding and recoding phase behavior of TDP43 reveals that phase separation is not required for splicing function

**DOI:** 10.1101/548339

**Authors:** Hermann Broder Schmidt, Ariana Barreau, Rajat Rohatgi

## Abstract

Intrinsically disordered regions (IDRs) often are fast evolving protein domains of low sequence complexity that can drive phase transitions and are commonly found in many proteins associated with neurodegenerative diseases, including the RNA processing factor TDP43. Yet, how phase separation contributes to the physiological functions of TDP43 in cells remains enigmatic. We combined systematic mutagenesis guided by evolutionary sequence analysis with a live-cell reporter assay of TDP43 phase dynamics to identify regularly-spaced hydrophobic motifs separated by flexible, hydrophilic segments in the IDR as a key determinant of TDP43 phase properties. This heuristic framework allowed us to customize the material properties of TDP43 condensates to determine effects on splicing function. Remarkably, mutants with increased or decreased phase dynamics, and even mutants that failed to phase separate, could mediate the splicing of a quantitative, single-cell splicing reporter and endogenous targets. We conclude that phase separation is not required for the function of TDP43 in splicing.

## Introduction

Trans-activating response (TAR) element DNA-binding protein of 43kDa (TDP43) is a regulator of RNA processing and found to be aggregated in many neurodegenerative diseases, including amyotrophic lateral sclerosis (ALS), frontotemporal dementia (FTD) and Alzheimer’s disease (Amador-Ortiz et al., 2007; Arai et al., 2006; Neumann et al., 2006). One of the best-defined functions of TDP43 is its role in skipping cryptic exons during alternative splicing to ensure that they do not disrupt the final mRNA message (Ling et al., 2015). TDP43 predominantly localizes to the nucleus, where it can exist both in a soluble form and in association with various nuclear bodies (Wang et al., 2002) such as paraspeckles (Naganuma et al., 2012; West et al., 2016), which are membrane-less protein-RNA condensates thought to form via liquid-liquid phase separation (LLPS) (Fox et al., 2018). When cells face stress, TDP43 redistributes to the cytoplasm and associates with stress granules (Colombrita et al., 2009; Dewey et al., 2011), another prominent type of liquid-like protein-RNA condensate (Protter and Parker, 2016).

TDP43 is comprised of a folded N-terminal domain (NTD) (Afroz et al., 2017; Mompean et al., 2016), two folded RNA-binding RRM domains (Lukavsky et al., 2013) and a low-complexity C-terminal domain (CTD) that is mostly disordered (Conicella et al., 2016; Jiang et al., 2013; Lim et al., 2016) (Fig. 1A). The CTD is a critical driver of TDP43 phase transitions (Conicella et al., 2016; Lim et al., 2016; Schmidt and Rohatgi, 2016) and is required for the recruitment of TDP43 to stress granules (Bentmann et al., 2012; Colombrita et al., 2009; Dewey et al., 2011) and presumably also to nuclear bodies (Fox et al., 2018; Hennig et al., 2015). Remarkably, the CTD harbors nearly all known human mutations in TDP43 that cause ALS (Buratti, 2015). Since some of these mutations have been shown to alter the material properties of TDP43 condensates (Gopal et al., 2017; Schmidt and Rohatgi, 2016) and to interfere with TDP43-mediated splicing (Arnold et al., 2013), it has been suggested that TDP43 phase separation is required for the function of TDP43 in RNA processing.

**Figure 1.**
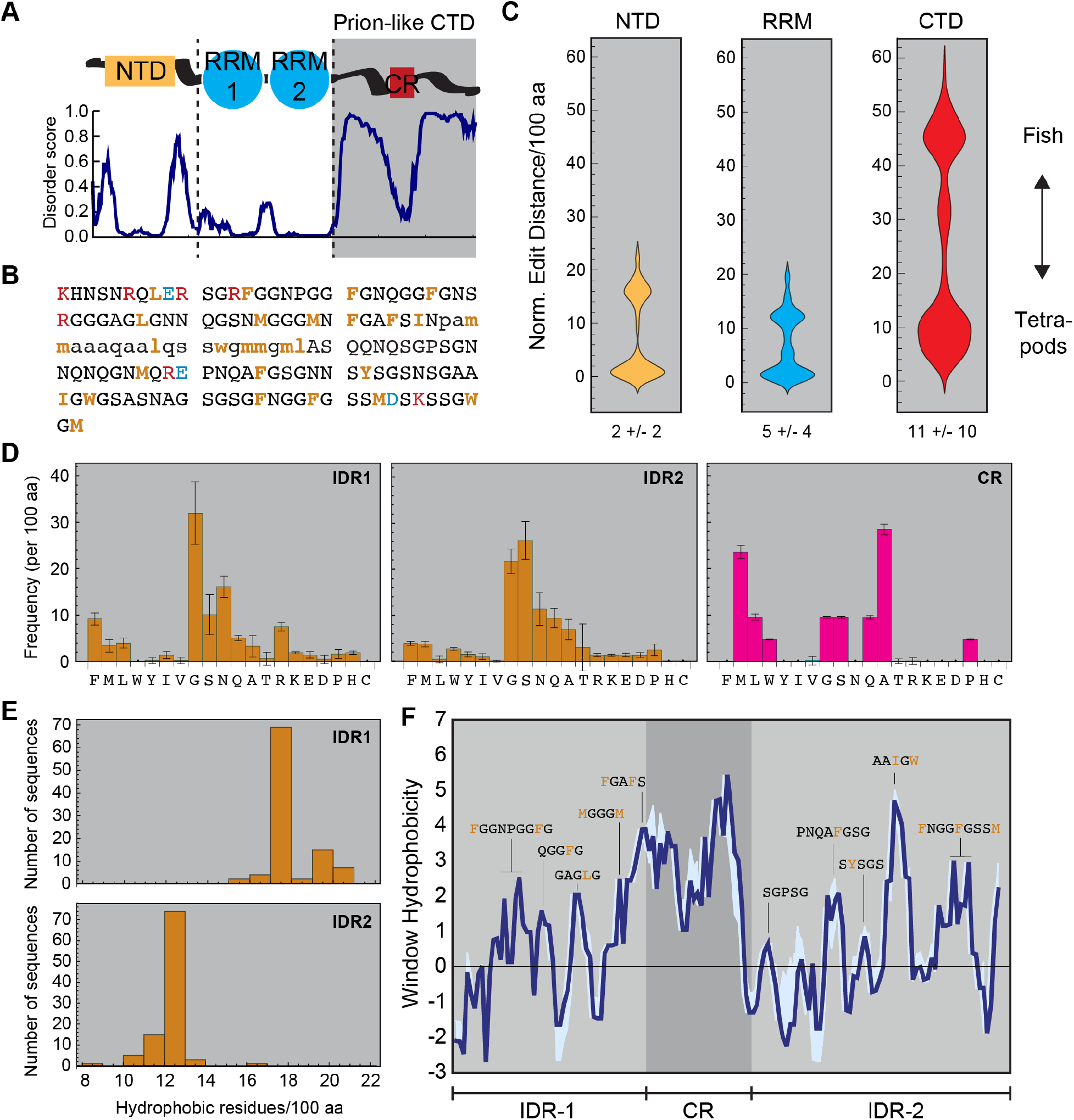
Identification of conserved sequence features in the TDP43 CTD. (**A**) TDP43 domain overview and disorder prediction. TDP43 is comprised of a folded N-terminal domain (NTD), two folded folded RNA-recognition motifs (RRM 1&2) and disorder-containing C-terminal domain (CTD) featuring a highly conserved region (CR). Disorder predictions are based on DisoPred (bioinf.cs.ucl.ac.uk/psipred) analysis for human TDP43 (UniProtKB Q13148). (**B**) Primary amino acid sequence of the human TDP43 CTD. Disordered regions are in capital and the CR in lower-case letters. Hydrophobic motifs are highlighted in bold orange, negatively-charged residues in blue and positively-charged residues in red. (**C**) Violin plots of the sequence divergence (represented as normalized Levenshtein distance per 100 aa) between the NTDs, RRMs and CTDs of 99 vertebrate TDP43 homologs. The plots show two clusters each, corresponding to fish and tetrapod TDP43 homologs. The numbers below the plots represent the median sequence changes in the entire dataset for the tetrapod cluster only (in parenthesis). (**D**) Amino acid composition of the intrinsically disordered regions (IDRs) and conserved regions (CRs) that comprise the TDP43 CTD. (**E**) Density distribution of the conserved hydrophobic residues (V, L, I, M, F, Y, W) in the IDRs of the TDP43 CTD. (**F**) Comparison of the hydrophobicity distribution across the vertebrate TDP43 CTD dataset. Hydrophobicity was calculated in a sliding window of 5 amino acids on the scale of Fauchère and Pliska (Fauchere and Pliska, 1983) (black trace: human CTD; light blue area: deviation amongst CTD homologs; sequence snippets: major hydrophobic windows).

We sought to test this model by exploring the evolutionary sequence space of vertebrate TDP43 CTDs and systematically determining the effect of sequence on both the material properties of TDP43 condensates and on the function of TDP43 in splicing. We find that aromatic residues in the TDP43 CTD are key drivers of phase separation and contribute to the recruitment of TDP43 to nuclear bodies. Furthermore, periodic (aromatic and non-aromatic) hydrophobic motifs embedded in a more hydrophilic sequence context determine the material properties of TDP43 condensates. Disease-causing or engineered single amino acid exchanges that alter this pattern interfere with TDP43 phase dynamics in predictable ways. Notably, neither mutations that increase or decrease the dynamics of TDP43 condensates, nor mutations that abrogated TDP43 phase transitions altogether, prevented the TDP43-dependent splicing of endogenous targets or a single-cell quantitative splicing assay. These findings suggest that condensate formation is not required for the function of TDP43 in exon skipping.

## Results

### The primary amino acid sequence of the TDP43 CTD is poorly conserved in vertebrates

The disorder-containing CTD (Fig. 1B) drives TDP43 phase separation and is a hotspot for mutations that cause ALS. These mutations can both alter the material properties of TDP43 condensates (Gopal et al., 2017; Schmidt and Rohatgi, 2016) and disrupt TDP43-mediated splicing (Arnold et al., 2013), suggesting that TDP43 condensation may be required for TDP43-dependent exon skipping. We sought to test this model by engineering TDP43 phases with custom material properties and measuring the consequences on TDP43 function in splicing. However, in order to customize TDP43 phase properties, we first had to decode the rules that relate TDP43 phase behavior to its amino acid sequence. Therefore, we turned to evolutionary sequence analysis and compiled a database with protein sequences of 99 vertebrate TDP43 homologs, which we split up into separate collections of NTDs, RRM domains and CTDs (Fig. S1A). As a simple measure of the sequence diversity within these collections, we calculated the number of amino acid exchanges, additions or subtractions (i.e. the Levenshtein or edit distance) normalized to a sequence length of 100 amino acids between all members of each category (Fig. 1C). As expected for low-complexity domains, the edit distance in the CTD collection is much higher than in the RRM and NTD collections (median 11 +/- 10 compared to 5 +/- 4 and 2 +/- 2, respectively), indicating that there is less selection pressure on the precise sequence of amino acids in the CTD. The sequences in each collection fall into either of two clusters (explaining the large median deviations), which correspond to fish and tetrapod TDP43 homologs. Within the tetrapod cluster, NTDs differ only in a single residue while CTDs vary by almost ten amino acids (Fig. 1C). Paradoxically, even though amino acid substitutions in the CTD have been frequent during vertebrate evolution, point mutations that lead to single amino acid changes and cause familial ALS in humans are most frequently found in the CTD (Kabashi et al., 2008; Sreedharan et al., 2008), highlighting the importance of decoding the sequence rules underlying TDP43 phase behavior in physiology and disease.

### Overall sequence composition and hydrophobic spacing are evolutionary conserved in the TDP43 CTD

Given the poor conservation of the primary amino acid sequence in the TDP43 CTD, we searched for higher-order sequence patterns that might be conserved. The TDP43 CTD is discontinuous and interrupted by a short, conserved region (CR) prone to adopt an α-helical fold, especially when TDP43 undergoes phase separation (Conicella et al., 2016; Schmidt and Rohatgi, 2016). This unusual (and complicating) arrangement (Fig. 1A) prompted us to separately analyze the amino-acid composition of the CR and each of the two intrinsically disordered regions (IDRs) that bracket the CR (Fig. S1A). There was a clear compositional bias in the IDRs and CRs compared to structured proteins in the Protein Data Bank (PDB), in particular the depletion of charged residues in both types of subdomains (Fig. 1D & Fig. S1B). Moreover, the IDRs were especially enriched in glycine residues (∼31% in IDR1 and ∼21% in IDR2), while alanine (∼28%) and methionine (∼24%) were abundant in the CR (Fig. 1D & Fig. S1B). This agrees with the reported helix propensity of these residues, which is lowest for glycine and high for methionine and alanine (Pace and Scholtz, 1998).

Beyond overall sequence composition, we noticed that hydrophobic residues (V, L, I, M, F, Y, W) occur in conserved intervals (Fig. 1B and Supplemental File 1). Indeed, both the number of hydrophobic residues (∼18 +/- 1 per 100 amino acids in IDR1 and ∼12 +/- 1 in IDR2) (Fig. 1E), as well as their relative position and spacing (Supplemental Files 1 & 2) are strikingly conserved in the IDRs. Because the spacer sequences between the hydrophobic motifs are rich in small and polar amino acids (Fig. 1B), the TDP43 IDRs can be characterized as a regular alternating pattern of hydrophobic and hydrophilic segments. To illustrate this periodic hydrophobic pattern, we calculated the hydrophobicity throughout the human TDP43 CTD in a sliding window of five amino acids using sixteen different hydrophobicity scales (Fig. S1C). Applying this analysis to our entire set of vertebrate TDP43 CTDs revealed that both the periodicity and the amplitude (i.e. the window hydrophobicity, hereafter shown on the scale of Fauchère and Pliska (Fauchere and Pliska, 1983)) of these motifs are well conserved (Fig. 1F). These findings suggest that the rules governing TDP43 phase separation and dynamics may be encoded in this conserved hydrophobic pattern.

### A strategy to rapidly assess the effects of changing CTD sequence on TDP43 phase behavior in living cells

We sought to measure how changing the periodic hydrophobic motifs in the TDP43 IDRs alters phase behavior in the physiological environment of living cells (rather than measuring phase behavior of the isolated protein *in vitro*). However, as we have demonstrated previously (Schmidt and Rohatgi, 2016), the small, diffraction-limited condensates formed by endogenous or fluorescently-labeled full-length TDP43 cannot be analyzed by live-cell imaging methods required to quantitatively characterize phase properties, such as fluorescence recovery after photobleaching (FRAP). To overcome this barrier, we replaced the RNA-binding RRM domains of TDP43 with GFP (Fig. S2A) to generate a reporter protein (hereafter called TDP43_RRM-GFP_) that assembles into micron-sized liquid droplets (Fig. 2A). These TDP43_RRM-GFP_ condensates are amenable to quantitative dynamic analysis in live cells and their phase properties are sensitive to single amino acid changes in the CTD, including changes that are known to cause ALS (Schmidt and Rohatgi, 2016). Importantly, the phase behavior of TDP43_RRM-GFP_ droplets resembles that of TDP43 condensates *in vitro* (Conicella et al., 2016; McGurk et al., 2018a; Molliex et al., 2015) and in neurons (Gopal et al., 2017). TDP43_RRM-GFP_ condensates are hence ideally suited for rapid, quantitative in-cell analysis of CTD sequence variants, allowing us to screen a panel of CTD mutants in which we systematically altered the composition and spacing of selected hydrophobic motifs or residues. Notably, we left the CR untouched in our mutagenesis studies since this region has been previously implicated in mediating interactions with other splicing factors such as hnRNPA2 (D’Ambrogio et al., 2009), making it difficult to deconvolve effects on protein interactions from effects on phase separation. We used the TDP43_RRM-GFP_ reporter assay to identify mutations in the CTD that lead to alterations in phase separation propensity or phase properties and then introduced these mutations in the context of full-length TDP43 to study their effect on splicing.

**Figure 2.**
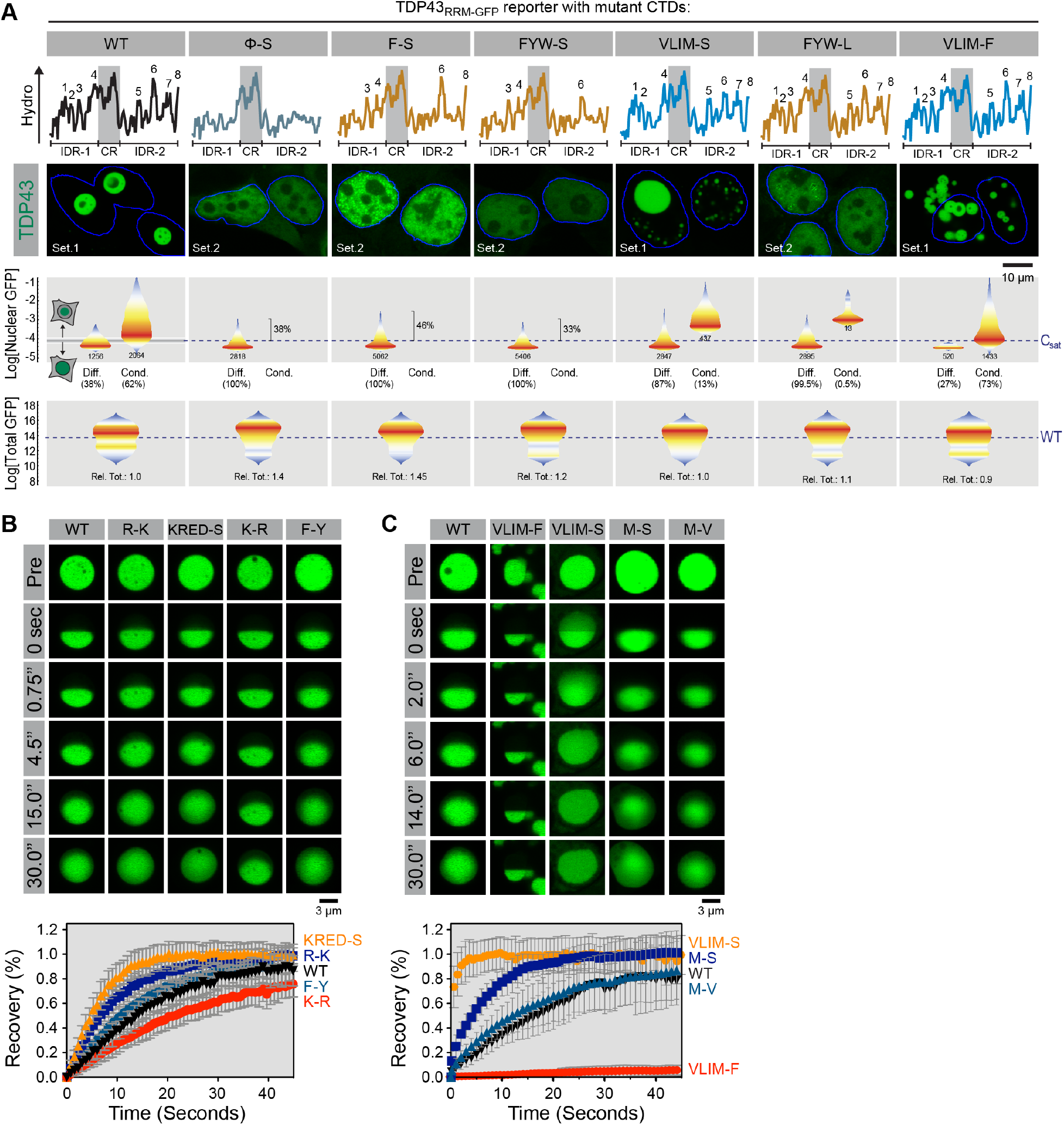
Aromatic residues and π-interactions are crucial drivers of TDP43 phase separation. (**A**) TDP43_RRM-GFP_ reporter constructs carrying the indicated mutations were transiently transfected into HEK-293T cells and analyzed by high-throughput automated confocal microscopy and flow cytometry. ***Top row:*** hydrophobicity plots for each CTD, with numbers highlighting the major hydrophobic clusters. ***Middle row:*** representative confocal images (63x) of cells transfected with various TDP43_RRM-GFP_ constructs. Blue outline demarcates nuclei; Set1 and Set2 indicate two different sets of photomultiplier settings used for optimal display only. ***Bottom rows:*** quantification of nuclear TDP43_RRM-GFP_ levels in cells with (‘Cond’) or without (‘Diff’) condensates measured by confocal images (20X) and total TDP43_RRM-GFP_ levels measured by flow cytometry. Data is presented as violin plots, with color-coding representing the probability density of cells in a particular fluorescence bin. Numbers and percentages below each graph reflect the absolute and relative number of transfected cells with (Cond) or without (Diff) TDP43_RRM-GFP_ condensates, respectively. For cells without condensates, percentages within in each graph reflect the number of cells with nuclear TDP43 fluorescence above the estimated critical concentration for phase separation. For total TDP43_RRM-GFP_ levels (bottom row), “Rel. Tot.” denotes the overall median cellular abundance of each CTD variant relative to the WT (n ≥ 10,000 cells). Dashed blue lines indicate the C_sat_ of WT TDP43_RRM-GFP_ (C_sat_) or the median total abundance of WT TDP43_RRM-GFP_ (WT), respectively. (**B**) and (**C**) Half-bleach FRAP experiments to determine the impact of the indicated CTD mutations on the dynamics of TDP43_RRM-GFP_ droplets in transiently-transfected HEK-293T cells. Plots show time-dependent, normalized fluorescence recovery. Plot markers represent mean values and error bars standard deviation (n ≥ 7).

### Aromatic residues in the TDP43 IDRs drive phase separation

Phase separation of TDP43_RRM-GFP_ was completely abrogated (Fig. 2A) when we eliminated the hydrophobic motifs in the IDR by changing all hydrophobic residues (VLIMFYW, hereafter abbreviated as Φ) to serine (all mutant CTD sequences used in this study are listed in Supplemental File 2). However, the Φ-S mutation also led to the partial mis-localization of TDP43_RRM-GFP_ to the cytoplasm, raising the possibility that the abundance of nuclear TDP43_RRM-GFP_ fell below the critical concentration required for phase separation (C_sat_). To estimate changes in C_sat_ caused by IDR mutations, we combined high-throughput automated confocal microscopy and flow cytometry to monitor the nuclear and total GFP fluorescence in thousands of cells expressing TDP43_RRM-GFP_ with wild-type (WT) or mutant IDRs (Fig. 2A). For a population of cells transfected with WT TDP43_RRM-GFP_, comparing nuclear GFP fluorescence in cells with TDP43_RRM-GFP_ droplets to cells with diffuse TDP43_RRM-GFP_ allowed us to estimate the C_sat_ of WT TDP43_RRM-GFP_, which was equal to ≈1/3 of the average expression level in the whole population (or ≈5 µM (Schmidt and Rohatgi, 2016)) (Fig. 2A). The nuclear abundance of the Φ-S mutant exceeded this threshold concentration in >38% of the cells, yet these cells did not show any signs of droplet formation (Fig. 2A). Hence, we conclude that Φ residues are crucial for TDP43 phase separation.

To parse the roles played by the different types of Φ residues, we first mutated all phenylalanine residues, the most dominant hydrophobic side chain in the TDP43 IDR (Fig. 1D), to serine (F-S). Even though the nuclear abundance of the F-S mutant was above the LLPS threshold in >46% of cells, it failed to form droplets (Fig. 2A). Phenylalanine could contribute to phase separation either because of its hydrophobicity, as proposed in the case of nucleoporin FG repeats (Labokha et al., 2013), or because of its aromatic structure and consequent ability to participate in π-interactions (Vernon et al., 2018). To assess the relative importance of hydrophobicity or aromaticity, we mutated either all aromatic or all aliphatic residues in the CTD to serine (FYW-S and VLIM-S, respectively). While the FYW-S mutant did not form TDP43_RRM-GFP_ droplets, even in cells where its concentration exceeded the LLPS threshold (Fig. 2A), the VLIM-S mutant could phase separate, although its C_sat_ was significantly elevated (Fig. 2A).

Given that there are 11 aromatic residues but only 9 aliphatic residues in the TDP43 IDRs, we also converted either all aromatics to leucine (FYW-L) or all aliphatics to phenylalanine (VLIM-F) in order to ensure that changes in phase separation were not merely due to a reduction of overall CTD hydrophobicity. Remarkably, we observed severe phase separation defects in the FYW-L mutant, despite the fact that the overall CTD hydrophobicity was only modestly altered (Fig. 2A). In contrast, the VLIM-F mutant formed clumps of small spherical assemblies at nuclear levels considerably below the C_sat_ for WT TDP43_RRM-GFP_ (Fig. 2A). Together, our findings suggest that TDP43 phase transitions in cells require aromatic residues, as *in vitro* (Li et al., 2018). Purely hydrophobic interactions, such as those seen in tropoelastin (Yeo et al., 2011), are not sufficient.

### TDP43_RRM-GFP_ phase separation is largely driven by π-π interactions

In addition to hydrophobic interactions, aromatic residues can also engage in π-π and cation-π interactions. For instance, cation-π interactions, especially involving arginine and tyrosine residues, govern the phase separation of FUS, an RNA-binding protein related to TDP43 (Bogaert et al., 2018; Qamar et al., 2018; Vernon et al., 2018; Wang et al., 2018). To test the requirement of cation-π interactions for TDP43 phase separation, we mutated all arginine residues to lysine (R-K) or all lysine residues to arginine (K-R). Both mutants formed droplets that were morphologically indistinguishable from WT TDP43_RRM-GFP_ droplets (Fig. 2B). Also, mutating all charged residues in the TDP43 IDR to serine (KRED-S) failed to abolish phase separation (Fig. 2B). We conclude that neither electrostatic nor arginine-mediated interactions are essential for phase separation of TDP43_RRM-GFP_ in cells.

While certain classes of interactions may have little effect on the formation of droplets *per se*, they could still influence the material properties of these droplets. Material properties of the large TDP43_RRM-GFP_ droplets can be readily assessed using half-bleach FRAP experiments (Schmidt and Rohatgi, 2016). While the WT and R-K droplets both displayed liquid-like dynamics, the R-K droplets were significantly more dynamic (Fig. 2B), suggesting that arginine-specific interactions (rather than charge-mediated electrostatic interactions) increase the viscosity of TDP43_RRM-GFP_ condensates. Arginine seems to be the most important charged residue for TDP43_RRM-GFP_ droplet dynamics because both the R-K and the KRED-S mutants increased droplet fluidity by a very similar degree (Fig. 2B). Consistent with this view, changing all lysines to arginines (K-R) significant reduced droplet dynamics (Fig.2B). Interestingly, mutating all phenylalanine residues to tyrosine (F-Y) did not alter TDP43 phase dynamics (Fig. 2B), suggesting that arginine does not discriminate between different aromatic side chains in TDP43, in contrast to observations in FUS (Bogaert et al., 2018; Qamar et al., 2018; Wang et al., 2018). Together, our data suggest that TDP43 phase transitions are mainly driven by π-π interactions involving aromatic residues, while arginine-mediated interactions influence the dynamics of the resulting condensates.

### Interactions involving aliphatic residues tune TDP43_RRM-GFP_ droplet properties

The aberrant morphology of VLIM-F condensates (Fig. 2A) suggested that an excess of aromatic residues can dramatically alter phase properties. Indeed, the VLIM-F mutant clumps did not behave like liquids— instead, the complete lack of fluorescence recovery in the bleached half (Fig. 2C) suggested a gel-like or solid-like character. In contrast, the VLIM-S mutant droplets were highly dynamic, almost instantaneously recovering fluorescence in the bleached half (Fig. 2C). Even the selective alteration of methionine residues to serine (M-S) increased TDP43_RRM-GFP_ droplet fluidity, whereas control methionine to valine (M-V) droplets displayed WT-like dynamics (Fig. 2C). Thus, although hydrophobicity is not sufficient to drive TDP43 phase transitions (as shown by the FYW-L mutant, Fig. 2A), it can have a dramatic influence on the phase properties of TDP43_RRM-_ GFP condensates. Moreover, our results suggest that a balance between aromatic and hydrophobic residues prevents solidification of TDP43 droplets and ensures that they maintain a liquid-like character. We propose that these requirements drive the conservation of hydrophobic motifs and overall sequence composition in the TDP43 CTD that we observed in our evolutionary sequence analysis (Fig. 1).

### The conserved spacing of hydrophobic motifs in the TDP43 IDRs regulates the liquid dynamics of TDP43_RRM-GFP_ droplets

An unexpected feature of our evolutionary analysis was the identification of conserved spacing between the hydrophobic motifs in the TDP43 CTD (Fig. 1E). In the CTD of human TDP43, both IDRs contain four major hydrophobic clusters each (Fig. 3A). By simply sliding the clusters together, we designed mutants where the total number of clusters was reduced to six, four or two (Figs. 3A & S2B). Importantly, the amino acid composition of the CTD and the overall hydrophobicity remained unaltered in these TDP43 variants.

**Figure 3.**
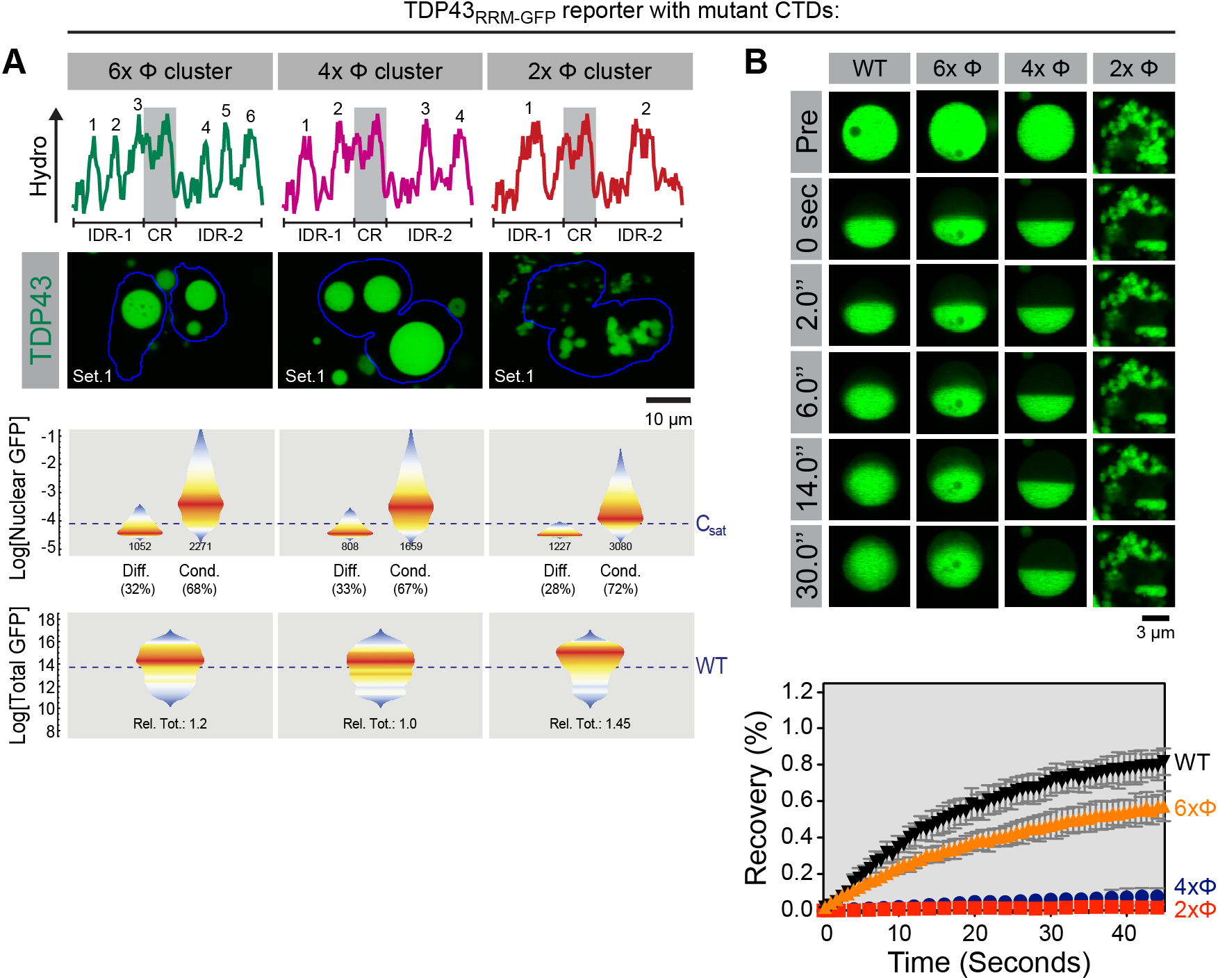
The spacing of hydrophobic residues across the CTD dictates the material properties of TDP43_RRM-GFP_ condensates. (**A**) Hydropathy plots (with numbers denoting hydrophobic peaks), cellular distribution, nuclear abundance and total cellular abundance of TDP43_RRM-GFP_ variants with altered spacing between hydrophobic peaks. Data is represented as in Fig. 2A (n ≥ 2,400 and 10,000 cells for nuclear and total GFP levels, respectively). See Fig. S2 for an illustration of how the mutants were constructed by sliding hydrophobic clusters. (**B**) Material properties of condensates formed by TDP43_RRM-GFP_ variants probed by half-bleach FRAP (as in Figs. 2B and 2C). Plot markers represent mean values and error bars standard deviation (n ≥ 5).

Whereas changing the number of hydrophobic (Φ) clusters from eight to six or to four did not change the morphology of TDP43_RRM-GFP_ reporter droplets, TDP43 variants with only two Φ clusters (2xΦ) formed amorphous assemblies that frequently mis-localized to the cytoplasm and behaved like solids in half-bleach experiments (Fig. 3A). Live cell imaging suggested that the 2xΦ mutant immediately aggregates after translation, as we observed a much earlier onset of phase transition (without a clearly detectable initial soluble state) and a greatly increased number of initial foci compared to WT (Fig. S2C). While the 6xΦ and 4xΦ mutants formed spherical droplets suggestive of a liquid-like genesis, the 6xΦ mutant had reduced dynamics and the 4xΦ droplets behaved like solids lacking any intra-droplet redistribution of TDP43_RRM-GFP_ (Fig. 3B). These observations support the model that the spacing between aromatic and aliphatic residues across the IDRs has an important influence on the material properties of TDP43_RRM-GFP_ phases by tuning intra-phase dynamics, preventing droplet solidification and aggregation.

Taken together, our mutagenesis of the TDP43 IDR revealed that evolutionarily conserved aromatic and aliphatic residues interrupted by hydrophilic and flexible segments are required for the formation of condensates with liquid-like properties. The importance of the periodic, interrupted nature of these clusters is highlighted by the fact that reducing the spacing between hydrophobic motifs clusters promotes aggregation and alters sub-cellular localization.

### The effect of point mutations on TDP43_RRM-GFP_ condensate dynamics can be predicted and rescued

To further test our periodic hydrophobicity model, we sought to determine if it could predict the effect of ALS-associated and engineered IDR mutations on the phase properties of TDP43_RRM-GFP_ droplets. We first focused on tryptophan 385, which is mutated to glycine in ALS patients (Millecamps et al., 2010) and forms the core of the major hydrophobic peak in IDR2 (Fig. 4A). The effects of the W385G mutation on TDP43 phase properties are not known. We expected that changing tryptophan 385 to glycine would enhance the internal dynamics of TDP43 condensates because it significantly reduces the hydrophobicity of IDR2 (Fig. 4A). This prediction conflicts with the prevalent view that disease mutations in RBPs promote the conversion of liquid phases into more solid-like assemblies (Lin et al., 2015; Molliex et al., 2015; Murakami et al., 2015; Patel et al., 2015). In accordance with our model, we observed faster fluorescence recovery kinetics in half-bleached W385G mutant TDP43 condensates (Fig. 4B, C). Thus, disease mutations in IDRs can also increase the liquidity of condensates, not just enhance their solid-like nature as commonly proposed.

**Figure 4.**
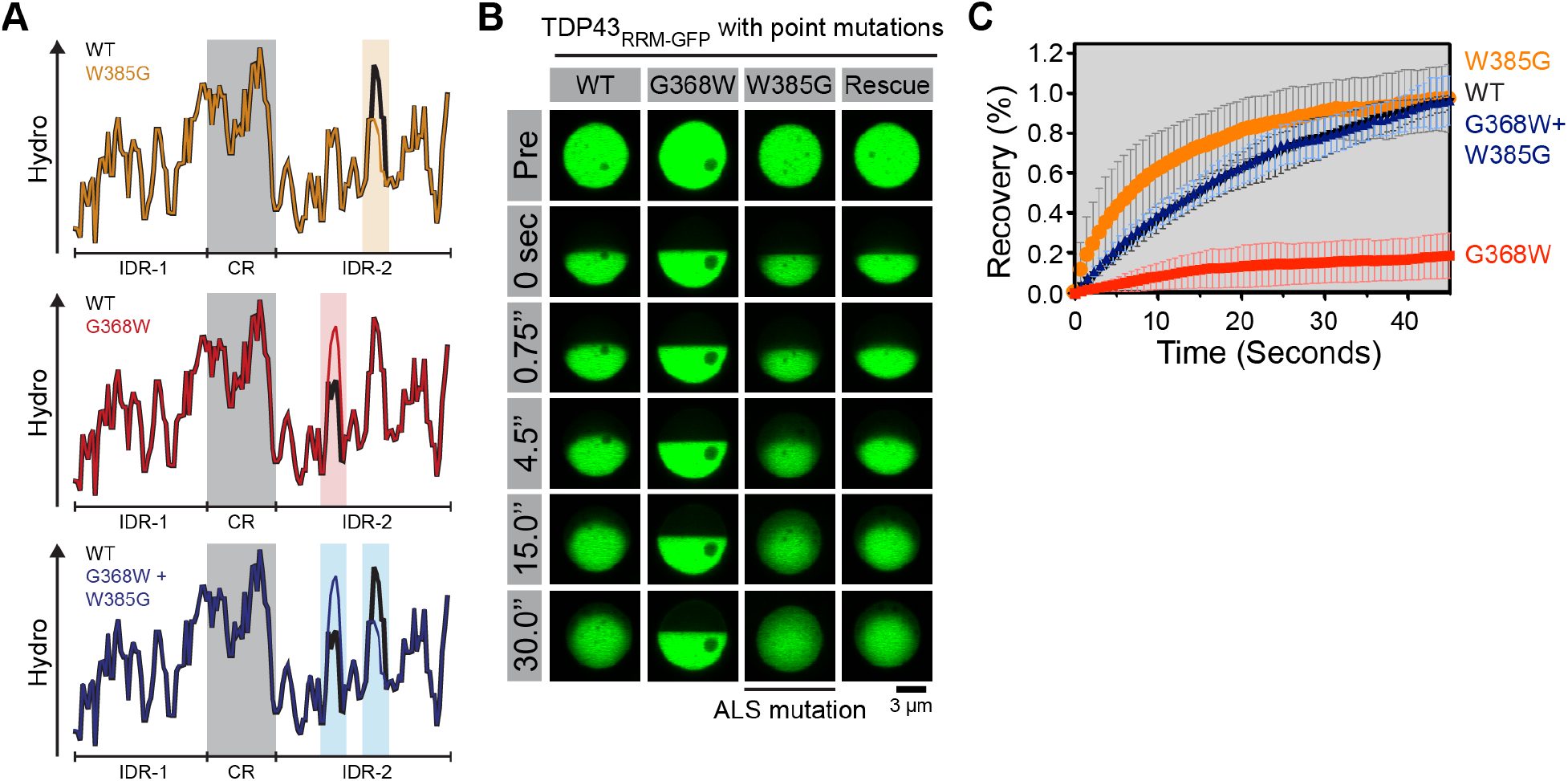
Compensatory second-site mutations can rescue altered phase dynamics caused by primary mutations. (**A**) Plots showing the effects of single and double point mutations on local hydrophobicity in the TDP43 CTD. (**B and C**) Material properties of TDP43_RRM-GFP_ droplets with single and double CTD mutations formed by transient transfection in HEK-293T cells. Plot markers represent mean values and error bars standard deviation (n ≥ 10).

Next, we studied another ALS-associated mutation, G348V (Buratti, 2015). This mutation increases the amplitude of a hydrophobic peak in IDR2 and, as expected, makes TDP43 phases less dynamic in half-bleach experiments (Fig. S3A, B). The effect of the G348V mutation was comparable to that of a synthetic G348F mutation (Fig. S3A, B), supporting the notion that hydrophobicity changes alone (without changes in aromaticity) can influence TDP43 phase properties. Similarly, increasing the height of a peak centered around glycine 309 by mutating it to phenylalanine, the most common hydrophobic residue found in the TDP43 IDR (Fig. 1D), decreased phase dynamics in agreement with our model (Fig. S3C, D). A control mutation of glycine 309 to serine, which did not change local hydrophobicity, had no effect on TDP43 phase dynamics (Fig. S3C, D).

As the most stringent test of our model that periodic hydrophobic motifs drive phase behavior, we aimed to rationally make second-site suppressor mutations that would “rescue” or restore the altered phase dynamics produced by a primary mutation. Guided by our model, we sought to reverse the fluidizing effect of the W385G mutation on TDP43_RRM-GFP_ condensates by increasing hydrophobicity at a distant site. We chose the minor hydrophobic peak around glycine 368, because mutating this residue to tryptophan would restore both the overall hydrophobicity and the amino-acid composition of the CTD (Fig. 4A). As expected, the elevated hydrophobicity produced by the G368W mutation in isolation makes TDP43 condensates markedly more solid (Fig. 4B, C). However, when the G368W mutation is made in combination with the W385G mutation, wild-type phase dynamics are restored in the double mutant (Fig. 4B, C). Taken together, our findings demonstrate that the intra-phase dynamics of TDP43 droplets can be tuned in predictable ways by single amino acid exchanges and suggest that the consequences of these mutations can be predicted based on how they influence local hydrophobicity.

### Aromatic residues drive the localization of full-length TDP43 to nuclear bodies

Our extensive screening of the CTD sequence space in our reporter system revealed that π-π interactions involving aromatic residues are the major driving force of TDP43 phase separation. To corroborate our findings in the context of full-length TDP43, we generated a clonal HEK-293T cell line carrying loss-of-function mutations in the *TARDBP* gene using Crispr/Cas9-mediated editing (Fig. S4) and then re-introduced GFP-tagged, full-length TDP43 variants by stable transduction (Fig. 5). In this assay, WT GFP-TDP43 consistently formed multiple nuclear foci in the majority of analyzed cells (≈70%) (Fig. 5), indicative of phase separation and localization to nuclear bodies (Maharana et al., 2018; Wang et al., 2002). Based on the literature, we expected that these foci correspond to paraspeckles (Naganuma et al., 2012; West et al., 2016), but they failed to co-localize with established markers of paraspeckles, nuclear speckles or PML bodies in HEK-293T cells (Fig. S5).

**Figure 5.**
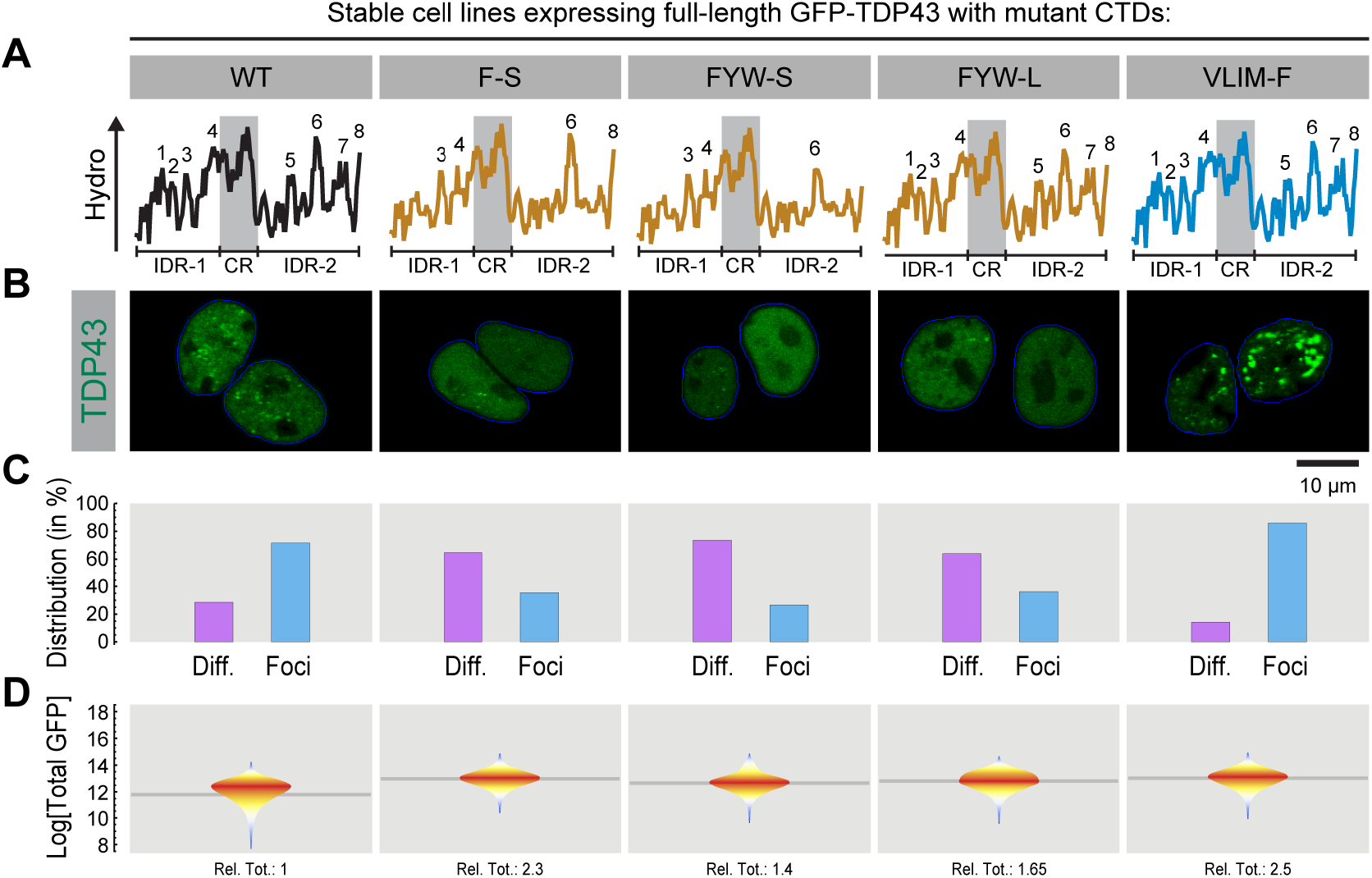
Aromatic residues in the CTD are important for the localization of full-length TDP43 to nuclear bodies. Hydropathy plots (**A**), nuclear distribution (**B**, **C**) and total cellular abundance (**D**) of full-length, GFP-tagged TDP43 (GFP-TDP43) variants with the indicated CTD mutations expressed by stable transduction in TDP43-/- HEK-293T cells. (**B**) Representative images of the nuclear distribution of each variant. (**C**) Percentage of cells showing nuclear foci (“Foci”) or a purely diffuse nuclear distribution (“Diff”); n ≥ 150 cells. (**D**) Distribution of total cellular GFP-TDP43 measured by flow cytometry and depicted as in Fig. 2A (n ≥ 150,000 cells). “Rel. Tot.” denotes the overall median cellular abundance of each GFP-TDP43 variant relative to the WT.

Both the number of cells with foci and the number of foci per cell were reduced in cells expressing GFP-TDP43 variants carrying the F-S, FYW-S or FYW-L mutant CTDs, despite being expressed at higher levels (Fig. 5). Notably, these mutations also significantly reduced the phase separation propensity of our TDP43_RRM-GFP_ reporter; Fig. 2A. In contrast, the VLIM-F mutation, which drove the solidification of TDP43_RRM-GFP_ droplets (Fig. 2A, C), led to the formation of larger, more numerous and clumped nuclear foci (Fig. 5). Together, these findings show that the phase behavior of the TDP43_RRM-GFP_ reporter correlates with the formation of nuclear bodies by full-length TDP43, likely because recruitment of TDP43 to nuclear bodies is driven by its phase separation (as has been previously shown for FUS (Hennig et al., 2015)).

### IDR-driven condensation is dispensable for TDP43-mediated exon skipping

Establishing the rules that govern TDP43 condensate formation and dynamics in cells put us in a position to systematically test the model that TDP43 function in RNA processing requires phase separation. We decided to focus on the role of TDP43 in mediating exon skipping during alternative splicing, because it (i) has been intensively studied (Ling et al., 2015; Polymenidou et al., 2011; Tollervey et al., 2011), (ii) is purportedly dependent on the CTD (Ayala et al., 2005; Buratti et al., 2005; D’Ambrogio et al., 2009), and (iii) is disrupted by ALS-associated mutations in the CTD (Arnold et al., 2013). In addition, it was recently demonstrated that splicing regulation by the low complexity domain-containing alternative splicing factor Rbfox depends on its phase separation (Ying et al., 2017).

To analyze the splicing activity of TDP43 variants rapidly, quantitatively and with single-cell resolution, we built a flow cytometry-compatible splicing reporter which uses a bidirectional promoter to co-express a splicing-sensitive fluorescent protein and full-length TDP43 (or variants thereof) from the same transiently-transfected plasmid (Fig. 6A). The splicing module is a full-length mEGFP fused to an mCherry construct interrupted by exon 9 of the CFTR gene (Buratti et al., 2001), which is skipped only in the presence of functional TDP43 (Figs. 6A and S6A). We established that the splicing of this module to express a functional mCherry was dependent on TDP43 by transfecting it into WT and TDP43-/- cells (Fig. S4). Skipping of exon 9 was severely reduced in TDP43-/- cells, assayed using either flow cytometry or reverse transcription-PCR (RT-PCR) to directly measure the distribution of splicing intermediates (Fig. S6). For our assays, this splicing module was co-expressed in TDP43-/- cells with full-length TDP43 CTD mutants carrying a BFP-P2A tag, which is post-translationally removed. To rigorously control our assay, the efficiency of TDP43-mediated exon skipping (mCherry fluorescence) was normalized both to the expression level of the splicing reporter (GFP fluorescence) as well to the expression level of the re-introduced TDP43 (BFP fluorescence) on a cell-by-cell basis (Fig. 6A).

**Figure 6.**
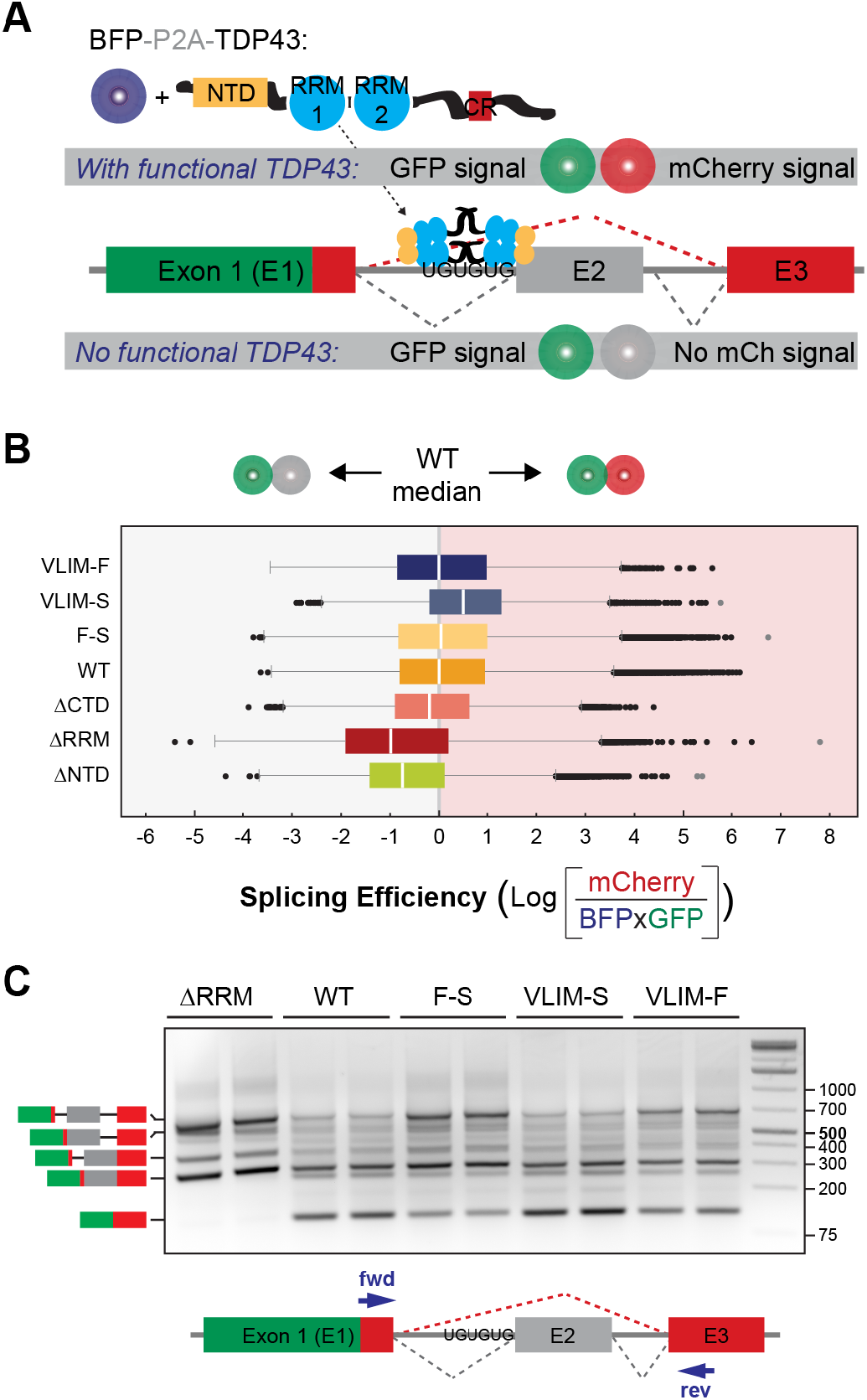
Mutations that alter phase separation can still support TDP43-mediated skipping of a model exon in a single-cell splicing reporter. (**A**) Splicing reporter design. Production of a dually-fluorescent GFP-mCherry fusion protein depends on the presence of functional full-length TDP43 to mediate skipping of exon 2 (“E2”, derived from the TDP43-regulated exon 9 of the CFTR gene). TDP43 and its CTD variants are expressed as fusions to BFP, which is post-translationally cleaved due to an intervening P2A site. The BFP-P2A-TDP43 and the GFP-mCherry splicing reporter are expressed from the same plasmid via a bi-directional promoter by transient transfection in TDP43-/- HEK- 293T cells. (**B**) Splicing efficiency of the indicated TDP43 compositional mutants quantitatively measured on a single-cell level by flow cytometry and expressed as the log of the mCherry/GFPxBFP ratio per cell. The grey vertical line indicates the median ratio for WT TDP43. (**C**) Exon skipping activity of TDP43 variants measured by RT-PCR using primers spanning exon 2 (see cartoon below the gel). Diagrams to the left denote the electrophoretic positions of the various splicing intermediates and products.

We first used this splicing reporter to compare the splicing efficiency of exogenous full-length TDP43 (WT) to TDP43 variants lacking the RNA-binding domains (ΔRRM), the N-terminal domain (ΔNTD) or the C-terminal domain (ΔCTD). Re-expression of TDP43WT in TDP43-/- cells rescued the exon skipping defect, whilst TDP43ΔRRM was non-functional (Fig. 6B, C). Both TDP43ΔNTD and TDP43ΔCTD showed splicing defects (Fig. 6C), as previously reported (Ayala et al., 2005; D’Ambrogio et al., 2009; Zhang et al., 2009).

We then tested the F-S CTD mutant, which failed to phase separate when introduced into the TDP43_RRM-GFP_ reporter (Fig. 2A) and reduced nuclear foci when introduced into full-length TDP43 (Fig. 5). This mutant was functional in our splicing assay, regardless of whether exon-skipping was assessed by flow cytometry (Fig. 6B) or RT-PCR (Fig. 6C). These unexpected observations suggested that phase separation is not required for the function of TDP43 in exon skipping.

To next address the question of whether changes in the dynamics of TDP43 phases can influence its splicing function, we tested the VLIM-S and VLIM-F CTD mutants, which make TDP43_RRM-GFP_ droplets markedly more fluid or more solid, respectively (Fig. 2). Despite the fact that these mutations change phase properties in opposite directions, both mutations have no detrimental effect on splicing (Fig. 6B, C).

Taken together, our data shows that the effects of mutations in the IDR on TDP43 phase behavior and splicing function are not correlated, thus supporting the conclusion that IDR-driven phase separation is dispensable for the function of TDP43 in exon skipping.

### TDP43 mutations that prevent phase separation do not block splicing of endogenous mRNAs

Given that our results disagreed with the view that phase separation plays a functional role in splicing, we sought to corroborate our results for three known, endogenous targets (*ATG4B, DNAJC5* and *ITPR3*) that contain cryptic exons (CEs) suppressed by TDP43 (Ling et al., 2015). The mRNAs encoded by these genes were analyzed by RT-PCR for the presence of the relevant cryptic exons (Fig. 7) in TDP43-/- cells (Fig. S4) or TDP43-/- cells stably expressing variants of full-length TDP43 (Fig. 5). As expected, disruption of TDP43 led to an increase in transcripts containing CEs at the expense of the fully spliced product (Fig. 7). Re-introduction of WT TDP43 restored the skipping of CEs in the mRNAs of all three targets (Fig. 7). Consistent with our results using the splicing reporter assay (Fig. 6), the F-S, FYW-S, FYW-L and VLIM-F mutants could all mediate CE skipping (Fig. 7). Taken together, these findings suggests that condensate formation is not required for TDP43-mediated exon skipping.

**Figure 7.**
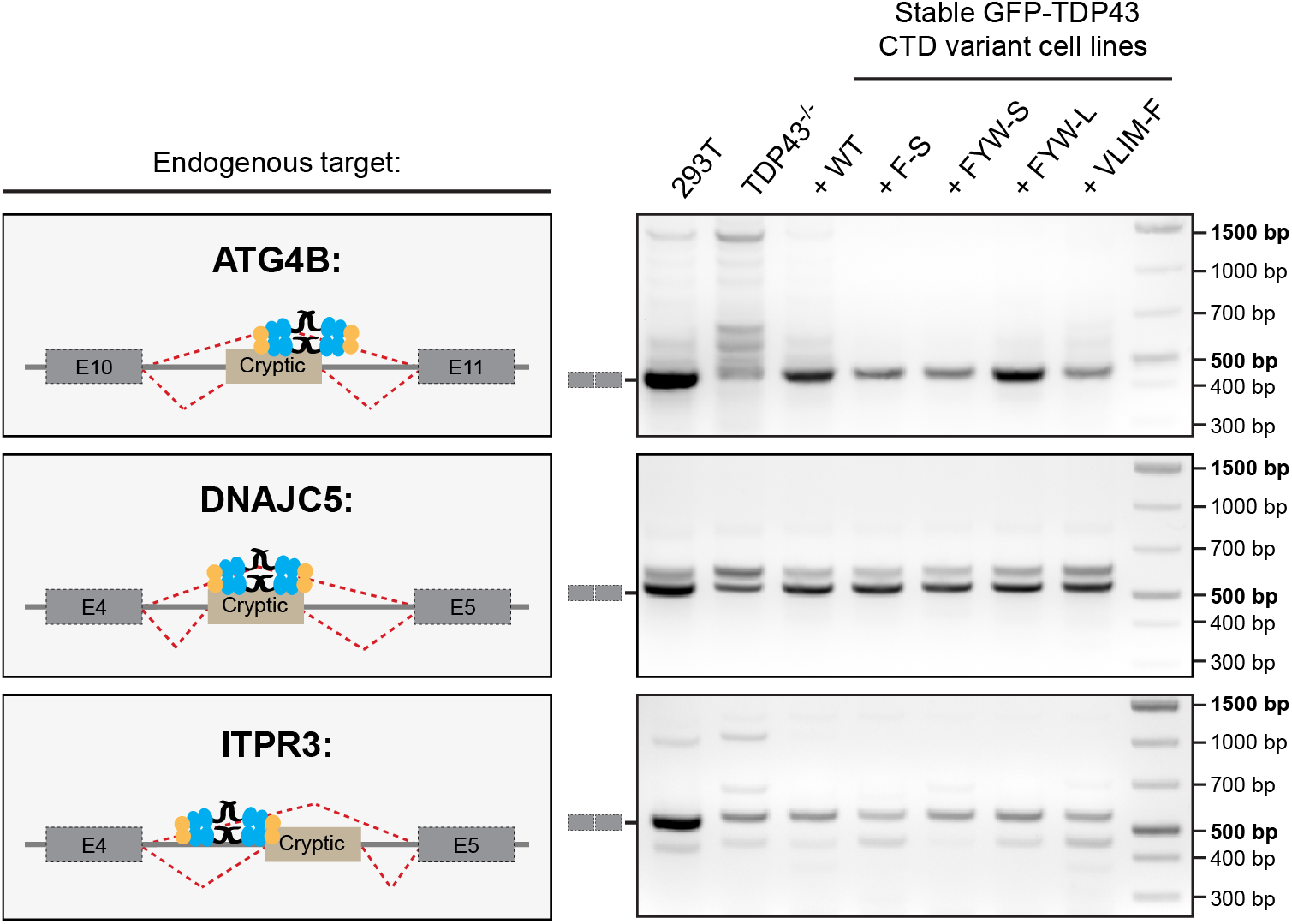
Mutations that alter phase separation can still support TDP43-mediated skipping of cryptic exons in endogenous genes. Splicing of cryptic exons in *ATG4B, DNAJC5* and *ITPR3* known to be regulated by TDP43 was assessed in WT, TDP43-/-, or TDP43-/- HEK-293T cells stably expressing the indicated variants of GFP-TDP43. See Fig. 5 for characterization of these cell lines, including assessment of total cellular GFP-TDP43 abundance by flow cytometry. Left cartoons show the position of the TDP43 binding site relative to the cryptic exon in each gene. Primers aligning to exons flanking the cryptic exon were used in RT-PCR (gel on right) to assess the relative abundance of the mRNA species that either retained or excluded the cryptic exon. The expected PCR products for the correctly spliced transcripts are marked to the left of the gels.

## Discussion

Phase transitions of proteins and nucleic acids into biomolecular condensates in cells are emerging as an important cellular mechanism to dynamically organize biochemical reactions in space and time (Alberti, 2017; Banani et al., 2017; Holehouse and Pappu, 2018). For example, proteins with the ability to phase separate are involved in all steps of gene expression (Malinovska et al., 2013), where LLPS has been suggested to play pivotal roles in chromatin organization (Larson et al., 2017; Strom et al., 2017), transcription (Hnisz et al., 2017) and alternative splicing (Ying et al., 2017). Many proteins that display phase separation behavior contain low-complexity regions such as the TDP43 CTD. While phase separation has been observed and phase behavior characterized for many proteins both *in vitro* and *in vivo*, the molecular code that relates amino acid sequence to phase behavior often remains enigmatic (Martin and Mittag, 2018). This has been a major barrier to understanding the functions of TDP43, an abundant RNA-binding protein that is mutated and aggregated in the neurodegenerative disease ALS (Johnson et al., 2009).

Using a systematic mutagenesis approach rooted in evolutionary sequence analysis, we uncovered that aromatic residues drive the phase separation of TDP43 (Figs. 2, 5), whereas a conserved sequence pattern dictates the material properties of the resultant TDP43 condensates (Figs. 3, 4). TDP43 contains two IDRs in its CTD, which both feature aromatic and aliphatic residues with a conserved spacing that are embedded in a flexible, polar sequence context (Fig. 1). The spacing of these ‘cohesive’ residues seems to be critical for maintaining TDP43 phase properties and providing a barrier to solidification (and ultimately aggregation), as mutations that reduce the spacing, while conserving overall IDR hydrophobicity, make TDP43 condensates more solid-like (Fig. 3).

In the case of IDR-mediated phase separation, significant uncertainty (and controversy) surrounds the identity of the structural motifs that mediate favorable cohesive interactions within condensates and hence determine their dynamics (or their liquid-like or solid-like character). Two models for such structural motifs include (1) multivalent interactions mediated by aromatic, charged or polar residues (Brady et al., 2017; Pak et al., 2016; Riback et al., 2017; Vernon et al., 2018) and (2) loosely adherent cross-β structures that lack extended hydrophobic segments to exclude water and to prevent the formation of insoluble amyloid aggregates (Hughes et al., 2018; Luo et al., 2018; Murray et al., 2017; Xiang et al., 2015). Examples of the latter are low-complexity, aromatic-rich, kinked segments (or LARKs) that have been proposed to form distorted β-sheets stabilized by inter-strand hydrogen bonds or aromatic stacking interactions (Guenther et al., 2018; Hughes et al., 2018; Luo et al., 2018).

Although the TDP43 IDRs contain aromatic residues (especially phenylalanine), only a few hydrophobic motifs in IDR1 are predicted to be LARK segments (Guenther et al., 2018; Hughes et al., 2018) (Fig. S7). This is in contrast to other RNA-binding proteins that form biomolecular condensates, such as FUS or hnRNPA1 (Lin et al., 2015; Molliex et al., 2015; Murakami et al., 2015; Patel et al., 2015), which are amongst the cellular proteins with the highest predicted LARK content (Hughes et al., 2018; Luo et al., 2018). It is also clear that mutations in IDR2, which is not predicted to contain any LARKs, can have dramatic effects on TDP43 phase dynamics (Fig. 4). While our sequence analysis and mutagenesis studies do not offer atomic insights to distinguish between these two structural models, our work demonstrates that regularly spaced π-π, hydrophobic and arginine-mediated interactions are determinants of TDP43 phase separation and behavior in cells. Thus, it appears that the CTD features multiple strategies to restrain TDP43 aggregation potential while maintaining the ability to phase separate, including optimized spacing of cohesive motifs, limiting LARK motifs and, as previously demonstrated, containing a conserved helical region (CR) that interrupts its IDR (Conicella et al., 2016).

Based on our mutagenesis studies, we developed a heuristic model to guide the design of mutations that change the dynamics of TDP43 phases in predictable ways (Fig. 4). Such systematic manipulation of the molecular code underlying TDP43 phase behavior should facilitate future investigations on how phase dynamics influence the various functions of TDP43 in physiological models. Surprisingly, we find that CTD-driven phase separation is dispensable for the function of TDP43 in exon skipping (Figs. 6, 7). The most parsimonious explanation for this observation is that ‘soluble’ TDP43 functions in splicing, whereas phase separation serves to buffer or restrict this splicing-competent pool. This model is consistent with prior work showing that the exon-skipping function of TDP43 is maintained when the CTD is replaced by a heterologous splicing-suppressor domain (Ling et al., 2015) and further supported by the recent observations that RNA-binding proteins are often prevented from phase separating in the nucleus by high concentrations of RNA (Banerjee et al., 2017; Maharana et al., 2018). Thus, strategies to solubilize TDP43, such as antisense oligonucleotides (Becker et al., 2017) or small molecule inhibitors (McGurk et al., 2018b; McGurk et al., 2018a) that prevent the accumulation and aggregation of TDP43, may also restore splicing defects in ALS patients. Despite a decade of work on this important protein involved in ALS, the central question of how phase separation and phase dynamics regulate the physiological function of TDP43 remains a challenge for the future.

## Materials and Methods

### Cell Lines

HEK-293T were obtained from ATCC and cultured at 37°C and 5% CO2 in high-glucose DMEM (GE Healthcare) supplemented with 10% FBS (Atlanta Biologicals), 2 mM L-glutamine (Gemini Biosciences), 1x MEM non-essential amino acids solution (Gibco), 40 U/ml penicillin and 40 µg/ml streptomycin (Gemini Biosciences).

To generate HEK-293T TDP43 knock-out cells, sgRNAs (Key Resources Table) targeting TDP43 were cloned into pX458 (Addgene #48138; (Ran et al., 2013)), transfected into HEK-293T cells using X-tremeGENE9 (Roche) and single-cell sorted for GFP-positive cells using a Sony SH800 flow cytometer. Individual clones were screened by immunoblotting using α-TDP43 (Proteintech #10782-2-AP) and α-alpha-tubulin (Sigma T6199) primary antibodies, and appropriate dye-conjugated secondary antibodies for detection with a LiCor Odyssey imager (see Key Resources Table). Single cell clones lacking TDP43 protein were further analyzed by qPCR after total RNA extraction with Trizol (Invitrogen), cDNA synthesis with the iScript kit (Bio-Rad) and quantification with the SYBR green kit (Bio-Rad) on a QuantStudio 5 real-time PCR system (Thermo Fisher Scientific). qPCR primers are listed in the Key Resources Table. HEK-293T TDP43 knock-out cells were cultured as the parent HEK-293T cells.

To generate the stable HEK-293T cell lines specified in the Key Resources Table, the indicated GFP-TDP43 CTD variants were cloned into pLenti-CMV Puro DEST (Addgene #17452; (Campeau et al., 2009)) and transfected into HEK-293T cells together with pMD2.G (Addgene #12259) and psPAX2 (Addgene #12260) to produce lentiviral particles for transduction. Virus was harvested 48 and 72 hours after transfection, pooled, filtered through non-binding 45µm syringe filters (Pall Corporation) and used to transduce HEK-293T TDP43 knock-out cells. After 24 hours, the virus-containing medium was removed and replaced with selection medium containing 2µg/ml Puromycin (Sigma-Aldrich). After 7 days of selection, cells were bulk sorted for similar GFP levels (where possible) using a Sony SH800 flow cytometer. All stable HEK-293T GFP-TDP43 cell lines were cultured as the parent HEK-293T cells.

### Bioinformatics analysis

A graphical overview of the analysis procedure is shown in Fig. S1 and source code available on Github (https://github.com/RohatgiLab/TDP43-analysis). In short, the NCBI blastp suite (Key Resources Table) was used to retrieve 394 vertebrate TDP43 homologs, with human TDP43 (UniProtKB Q13148) as the query in a search limited to taxid:7742. A species and isoform filter was applied to remove sequence redundancy, reducing the dataset to 101 TDP43 homologs. Based on sequence alignments to the well-characterized RNA binding domain of human TDP43 (PDB:4bs2, (Lukavsky et al., 2013)), the dataset was split up into sub-collections of N-terminal domains, RRM domains and C-terminal domains. The latter was further dissected into the IDR and CR collections using sequence alignments to the previously described CR (Conicella et al., 2016; Schmidt and Rohatgi, 2016). To determine the density distribution of hydrophobic residues in the IDRs (but not CRs), we calculated the number of V, L, I, M, F, Y and W residues in each IDR and normalized the values to a reference length of 100 amino acids. Hydrophobicity was calculated according to the scale of (Fauchere and Pliska, 1983), which we re-scaled to range between 0 and 1 for clarity (as in (Schmidt and Goerlich, 2015)). For the comparison of hydrophobicity scales in Supplemental Figure 1C, the ProtScale tool from the ExPASy suite was used (see Key Resources Table).

The following published dataset was used for comparison: a compilation of folded proteins from the PDB (Schmidt and Goerlich, 2015).

### DNA constructs

All plasmids used in this study are summarized in the Key Resources Table. Sequences and maps are made available on Addgene. All compositional IDR mutants were designed *in silico*, synthesized as gBlocks by Integrated DNA Technologies and introduced into TDP43 using Gibson cloning. IDR point mutations were introduced by Gibson-based site-directed mutagenesis.

### Phase separation reporter assay

HEK-293T cells were seeded onto acid-washed coverslips coated with 0.1% gelatin (Sigma-Aldrich) in 24-well plates at 5×10^4^ cells/well, allowed to adhere overnight and transfected with the indicated reporter constructs (500 ng/well) using X-tremeGENE 9 (Roche). 24 hours after transfection, cells were fixed with 4% PFA for 15 minutes at room temperature, washed 3x with PBS and permeabilized with 0.1% Triton X-100 (Sigma-Aldrich) in PBS for 15 minutes at room temperature. Cells were then washed 3x with PBS and mounted onto glass slides using ProLong Diamond mounting medium containing DAPI (Molecular Probes). After curing over night at room temperature, z-stacks were automatically taken for all samples at 16 randomly-selected positions across each coverslip with a Leica DMI-6000B microscope equipped with a Yokogawa CSU-W1 spinning disk confocal unit at 20x resolution. A 488-nm laser was used for GFP excitation and an Andor iXon Ultra DU888 EMCCD camera for signal detection.

For quantification, custom-written Mathematica pipelines were used that are available on Github (https://github.com/RohatgiLab/TDP43-analysis). In short, the ImageFocusCombine function of Mathematica (Wolfram Research) was first used to make z-projections for each acquired z-stack. Using the DAPI channel, nuclear masks were then generated and used to (i) extract single nuclei and (ii) measure nuclear GFP levels. To subsequently compare the GFP levels of TDP43 droplet-free or droplet-containing nuclei, the morphological component analysis package of Mathematica was used to automatically detect spherical objects and classify the extracted nuclei accordingly. Mis-classifications where manually corrected if necessary. Non-transfected cells (within the population of transfected cells) were excluded from analysis by filtering based on the nuclear GFP signal of a deliberately non-transfected control cell population. At least 2,000 nuclei per sample were quantified.

To compare total reporter expression levels, HEK-293T cells were seeded in 24-well plates at 5×10^4^ cells/well, transfected as above, harvested 24 hours later and at least 10,000 analyzed by flow cytometry with a BD Accuri C6 instrument (excitation laser: 488 nm, emission filter: 586/40 nm).

### Phase dynamics reporter assay

HEK-293T cells were seeded into 8-well ibidi µ-slides (3 × 10^4^cells/well), allowed to adhere overnight and transfected with the indicated reporter constructs (300 ng/well) using X-tremeGENE 9 (Roche). 24 later, the culture medium was exchange with L-15 medium (Gibco) supplemented with 10% FBS, and cells imaged as previously described (Schmidt and Rohatgi, 2016) using a Leica SP8 laser-scanning confocal microscope equipped with a 488-nm laser, 63x glycerol objective (NA 1.3) and a temperature-controlled incubation chamber (Life Imaging Services) set to 37°C.

Half-bleach experiments were recorded and quantified using the FRAP module of the Leica Application Suite X software. To ensure that only similarly-sized droplets of comparable intensities were bleached, the same bleach window and detector settings were used. For each recorded time point (t), the fluorescence intensities within the bleached droplet hemisphere were then normalized to the fluorescence intensity of the corresponding unbleached droplet hemisphere. These normalized, time-dependent fluorescence intensities It were then used to calculate the fluorescence recovery (FR) according to the following formula: FR(t) = (I_t_ - I_t0_) / (Ibefore bleaching-I_t0_), with t0 being the first time point observed after photobleaching. At least five individual droplets from different cells were analyzed and GraphPad Prism was used to plot replicate measurements (mean ± SD). WT reporter constructs were included in all experiments as a control for comparison across dataset. For all measurements and initial fluorescence signal quantifications, investigators were blinded to prevent bias.

### Live cell imaging of TDP43 phase formation

To follow TDP43 reporter phase formation over time, cells were transfected as above and imaged in L15 medium (Gibco) supplemented with 10% FBS at 37°C every hour for a total of 24 hours after transfection with a Leica DMI-6000B microscope equipped with a Yokogawa CSU-W1 spinning disk confocal unit. A 488-nm laser was used for GFP excitation and an Andor iXon Ultra DU888 EMCCD camera for signal detection.

Onset of LLPS was determined by identifying the time point of foci formation, and the number of initial foci per cell in the corresponding frames were then counted using the morphological component detection function of Mathematica (Wolfram Research). The data was plotted and analyzed with GraphPad Prism using unpaired t-tests.

### Analysis of GFP-TDP43 foci in stable cell lines

Stable HEK-293T GFP-TDP43 CTD variant cell lines (see Experimental Model and Subject Details) were seeded into 8-well ibidi µ-slides (3 × 10^4^ cells/well), allowed to adhere overnight and imaged live in L15 medium (Gibco) supplemented with 10% FBS at 37°C using a Leica SP8 laser-scanning confocal microscope equipped with a 488-nm laser, 63x glycerol objective (NA 1.3) and a temperature-controlled incubation chamber (Life Imaging Services) set to 37°C. Nuclei were classified as foci-containing or foci-free using a combination of machine learning and manual classification in Mathematica (Wolfram Research). Source code is available at (https://github.com/RohatgiLab/TDP43-analysis).

For immunofluorescence staining, GFP-TDP43 CTD variants were seeded onto acid-washed coverslips coated with 0.1% gelatin (Sigma-Aldrich) in 24-well plates at 5×10^4^ cells/well, allowed to adhere overnight and fixed with 4% PFA for 15 minutes at room temperature. Cells were then washed 3x with PBS and treated with blocking buffer (PBS supplemented with 0.1% Triton X-100, 1% donkey serum and 10mg/ml BSA) for 30 minutes at room temperature. Incubation with primary antibodies against NONO (1:100), SFPQ (1:500), SC35 (1:500) and PML (1:500) (Key Resources Table) was performed overnight at 4°C in blocking buffer. After washing 3x with blocking buffer, cells were then incubated with fluorophore-conjugated secondary antibodies (Key Resources Table) for 2 hours at room temperature. Following three washes with PBS + 0.1% Triton X-100, samples were mounted using ProLong Diamond mounting medium containing DAPI (Molecular Probes) and imaged on a Leica SP8 laser-scanning confocal microscope using 488-nm, 561-nm and 633-nm excitation lasers and a 64x oil immersion objective (NA 1.4).

To compare total GFP-TDP43 levels, HEK-293T cells were seeded in 6-well plates and analyzed 48 hours later by flow cytometry with a BD Accuri C6 instrument (excitation laser: 488 nm, emission filter: 586/40 nm).

### Splicing reporter assays

HEK293T TDP43 knock-out cells were seeded into standard P6 tissue culture plates (at 4 × 10^5^ cells/well), allowed to adhere overnight and transfected with the indicated splicing reporter constructs (1 µg/well) using X-tremeGENE 9 (Roche). As shown in the Key Resources Table, each reporter comprised–expressed from a bidirectional promoter–the splicing module shown in Fig. 6A and a full-length TDP43 variant fused to an N-terminal, post-translationally cleaved P2A-BFP tag. 24 hours after transfection, cells were harvested by trypsinization for flow cytometry or using Trizol reagent (Ambion) for RNA extraction.

All flow cytometry-based splicing assays were analyzed with a Sony SH800 flow cytometer. Instruments parameter settings were adjusted following the compensation guidelines using cells either expressing BFP, GFP or mCherry, respectively. Per analysis run, data for ∼20,000 BFP-positive single cells was collected and analyzed with custom pipelines available on Github (https://github.com/RohatgiLab/TDP43-analysis) to calculate and plot the mCherry/(GFPxBFP) fluorescence intensity ratio on a single cell level. This ratio reflects the splicing efficiency of a given TDP43 variant in our assay, normalized to both the abundance of the splicing reporter (GFP signal) as well as the approximate TDP43 level (BFP signal) per cell. At least 73,000 measurements per variant from two independent biological replicates were analyzed and compared to WT using Mann-Whitney tests, showing that all medians were statistically different with p-values below 0.05.

To validate and corroborate the flow cytometry-based splicing measurements, total RNA was extracted using the Trizol reagent (Ambion) according to the manufacturer’s instructions, transcribed into cDNA using the iScript kit (Bio-Rad) and analyzed by analytical RT-PCR using the GoTaq Green kit (Promega) with primers 1171 and 1174 (see Key Resources Table). PCR products were separated by electrophoresis on 1.5-2% agarose gels and documented with an AlphaImager EC system (Alpha Innotech).

### Splicing assay for endogenous TDP43 target RNAs

To test for splicing defects in the previously reported endogenous TDP43 target RNAs ATG4B and ITPR3 (Ling et al., 2015), total RNA was first extracted from >1.0×10^6^cells using Trizol reagent (Ambion) following the manufacturer’s instructions, then transcribed into cDNA using the iScript kit (Bio-Rad) and subsequently analyzed using the GoTaq Green kit (Promega) for analytical RT-PCRs with the primers listed in the Key Resources Table. PCR products were electrophoresed on 1.5-2% agarose gels and documented with an AlphaImager EC system (Alpha Innotech).

## Acknowledgments

We thank Lindsay Becker for sharing antibodies, as well as Bede Portz and Steven Boeynaems for critical reading of the manuscript. We are grateful for support from the Deutsche Forschungsgemeinschaft (SCHM 3082/2-1 to H.B.S.), the National Institutes of Health (DP2 GM105448 & R35 GM118082 to R.R.) and a Stanford Undergraduate Advising and Research grant (to A.B.).

