## Supplemental Material for "Decoding and recoding phase behavior of TDP43 reveals that phase separation is not required for splicing function"

**KEY RESOURCES TABLE**

| REAGENT or RESOURCE | SOURCE | IDENTIFIER |
|---|---|---|
| **Antibodies** | | |
| Rabbit polyclonal anti-TDP43 | Proteintech | Cat# 10782-2-AP |
| Mouse monoclonal anti-NONO | Novus Biologicals | Cat# NBP2-02060 |
| Mouse monoclonal anti-SFPQ | Sigma-Aldrich | Cat# SAB4200501 |
| Mouse monoclonal anti-SC35 | Abcam | Cat# ab11826 |
| Rabbit monoclonal anti-PML | Abcam | Cat# ab179466 |
| Mouse monoclonal anti-α-Tubulin | Sigma-Aldrich | Cat# T6199 |
| Donkey anti-Rabbit IgG (H+L) Highly Cross-Adsorbed Secondary Antibody, Alexa Fluor 594 | Thermo Fisher Scientific | Cat# A-21207 |
| Donkey anti-Mouse IgG (H+L) Secondary Antibody, Alexa Fluor 647 | Thermo Fisher Scientific | Cat# A-31571 |
| IRDye 680LT Donkey anti-Mouse IgG (H+L) | Li-Cor | Cat# 926-68022 |
| IRDye 800CW Donkey anti-Rabbit IgG (H+L) | Li-Cor | Cat# 926-32213 |
| **Chemicals, Peptides, and Recombinant Proteins** | | |
| Trizol reagent | Ambion | Cat# 15596018 |
| Puromycin | Sigma-Aldrich | Cat#P8833 |
| **Critical Commercial Assays** | | |
| X-tremeGENE 9 | Roche | Cat# 06366236001 |
| iScript cDNA Synthesis Supermix | Bio-Rad | Cat# 1708891 |
| iTaq Universal SYBR Green Supermix | Bio-Rad | Cat# 1725120 |
| GoTaq® Green Master Mix | Promega | Cat# M7123 |
| **Deposited Data** | | |
| Vertebrate TDP43 homolog database | This study | https://github.com/RohatgiLab/TDP43-analysis |
| **Experimental Models: Cell Lines** | | |
| Human 293T cells | ATCC | CRL-3216 |
| Human 293T TDP43$^{-/-}$ cells | This study | N.A. |
| Human 293T TDP43$^{-/-}$ GFP-TDP43 CTD$^{WT}$ cells | This study | N.A. |
| Human 293T TDP43$^{-/-}$ GFP-TDP43 CTD$^{F-S}$ cells | This study | N.A. |
| Human 293T TDP43$^{-/-}$ GFP-TDP43 CTD$^{FYW-S}$ cells | This study | N.A. |
| Human 293T TDP43$^{-/-}$ GFP-TDP43 CTD$^{FYW-L}$ cells | This study | N.A. |
| Human 293T TDP43$^{-/-}$ GFP-TDP43 CTD$^{VLIM-F}$ cells | This study | N.A. |
| **Oligonucleotides** | | |
| 5'-caccgagatgctggctggggaaatc-3' | This study | 704 TARDBP-ko-1-fwd |
| 5'-aaacgatttccccagccagcatctc-3' | This study | 705 TARDBP-ko-1-rev |
| 5'-caccgaacatccgatttaatagtgt-3' | This study and (Shalem et al., 2014) | 706 TARDBP-ko-2-fwd |
| 5'-aaacacactattaaatcggatgttc-3' | This study and (Shalem et al., 2014) | 707 TARDBP-ko-2-rev |
| 5'-tggacgatggtgtgactgcaa-3' | (Prudencio et al., 2012) | 812 HsTARDBP qPCR fwd |
| 5'-agagaagaactcccgcagctca-3' | (Prudencio et al., 2012) | 813 HsTARDBP qPCR rev |
| 5'-gttcatgcgcttcaaggtg-3' | This study | 1171 Cryptic Exon fwd |

| | | |
|---|---|---|
| 5’-ttggtcaccttcagcttgg-3’ | This study | 1174 Cryptic Exon rev |
| 5’-ggaccagcgttccctgtgc-3’ | This study | 1019 HsATG4B fwd |
| 5’-caccaatcattgaagtcatc-3’ | This study | 1020 HsATG4B rev |
| 5’-tccttgggttggacaagaac-3’ | This study | 1311 HsDNAJC5 fwd |
| 5’-ttgaacccgtcagtgtggta-3’ | This study | 1312 HsDNAJC5 rev |
| 5’-gcggcgcagatggagcag-3’ | This study | 1021 HsITPR3 fwd |
| 5’-ccgaggtggcagactggc-3’ | This study | 1022 HsITPR3 rev |
| **Recombinant DNA** | | |
| pHBS838 TDP43$_{\text{RRM-GFP}}$ WT | (Schmidt and Rohatgi, 2016) | Addgene #98249 |
| pHBS941 TDP43$_{\text{RRM-GFP}}$ Φ-S | This study | Addgene #121992 |
| pHBS1383 TDP43$_{\text{RRM-GFP}}$ FYW-S | This study | Addgene #118791 |
| pHBS931 TDP43$_{\text{RRM-GFP}}$ F-S | This study | Addgene #107799 |
| pHBS1385 TDP43$_{\text{RRM-GFP}}$ VLIM-S | This study | Addgene #118792 |
| pHBS935 TDP43$_{\text{RRM-GFP}}$ M-S | This study | Addgene #107800 |
| pHBS1384 TDP43$_{\text{RRM-GFP}}$ FYW-L | This study | Addgene #118793 |
| pHBS1386 TDP43$_{\text{RRM-GFP}}$ VLIM-F | This study | Addgene #118794 |
| pHBS1037 TDP43$_{\text{RRM-GFP}}$ 6xΦ | This study | Addgene #107804 |
| pHBS1038 TDP43$_{\text{RRM-GFP}}$ 4xΦ | This study | Addgene #107805 |
| pHBS1039 TDP43$_{\text{RRM-GFP}}$ 2xΦ | This study | Addgene #107806 |
| pHBS934 TDP43$_{\text{RRM-GFP}}$ M-V | This study | Addgene #107801 |
| pHBS1387 TDP43$_{\text{RRM-GFP}}$ KRED-S | This study | Addgene #118795 |
| pHBS1052 TDP43$_{\text{RRM-GFP}}$ R-K | This study | Addgene #107856 |
| pHBS1388 TDP43$_{\text{RRM-GFP}}$ K-R | This study | Addgene #118796 |
| pHBS1256 TDP43$_{\text{RRM-GFP}}$ W385G | This study | Addgene #107807 |
| pHBS1291 TDP43$_{\text{RRM-GFP}}$ G368W | This study | Addgene #107837 |
| pHBS1292 TDP43$_{\text{RRM-GFP}}$ G368W+W385G | This study | Addgene #107838 |
| pHBS966 TDP43$_{\text{RRM-GFP}}$ G309F | This study | Addgene #107822 |
| pHBS968 TDP43$_{\text{RRM-GFP}}$ G309S | This study | Addgene #107824 |
| pHBS1280 TDP43$_{\text{RRM-GFP}}$ G348F | This study | Addgene #107830 |
| pHBS1279 TDP43$_{\text{RRM-GFP}}$ G348V | This study | Addgene #107831 |
| pHBS1147 GFP-TDP43 | This study | Addgene #118797 |
| pHBS1410 GFP-TDP43 F-S | This study | Addgene #118799 |
| pHBS1411 GFP-TDP43 FYW-S | This study | Addgene #118800 |
| pHBS1412 GFP-TDP43 FYW-L | This study | Addgene #118801 |
| pHBS1413 GFP-TDP43 VLIM-F | This study | Addgene #118802 |
| pHBS1389 IBB-GFP-mCherry3E | This study | Addgene #118803 |
| pHBS1396 [IBB-GFP-mCherry3E]-[BFP-P2A-TDP43 WT] | This study | Addgene #118804 |
| pHBS1393 [IBB-GFP-mCherry3E]-[BFP-P2A-TDP43 ΔRRM] | This study | Addgene #118805 |
| pHBS1394 [IBB-GFP-mCherry3E]-[BFP-P2A-TDP43 ΔCTD] | This study | Addgene #118806 |
| pHBS1392 [IBB-GFP-mCherry3E]-[BFP-P2A-TDP43 ΔNTD] | This study | Addgene #118807 |
| pHBS1401 [IBB-GFP-mCherry3E]-[BFP-P2A-TDP43 F-S] | This study | Addgene #118808 |
| pHBS1404 [IBB-GFP-mCherry 3E]-[BFP-P2A-TDP43 VLIM-S] | This study | Addgene #118809 |

| | | |
|---|---|---|
| pHBS1405 [IBB-GFP-mCherry 3E]-[BFP-P2A-TDP43 VLIM-F] | This study | Addgene #118810 |
| pHBS1398 [IBB-GFP-mCherry3E]-[BFP-P2A-TDP43 G368W] | This study | Addgene #118811 |
| pHBS1397 [IBB-GFP-mCherry3E]-[BFP-P2A-TDP43 W385G] | This study | Addgene #118812 |
| pHBS1399 [IBB-GFP-mCherry3E]-[BFP-P2A-TDP43 G368W+G385W] | This study | Addgene #118813 |
| pHBS1400 [IBB-GFP-mCherry3E]-[BFP-P2A-TDP43 G295F] | This study | Addgene #118814 |
| pHBS952 TARDBP43-Exon1-ko-sgRNA1-SpCas9 | This study | Addgene #107857 |
| pHBS953 TARDBP43-Exon2-ko-sgRNA2-SpCas9 | This study | Addgene #107858 |
| pMD2.G | Didier Trono Lab, EPFL, Switzerland | Addgene #12259 |
| psPAX2 | Didier Trono Lab, EPFL, Switzerland | Addgene #12260 |
| **Software and Algorithms** | | |
| blastp suite | NCBI | https://blast.ncbi.nlm.nih.gov/Blast.cgi |
| ProtScale | ExPASy | https://web.expasy.org/protscale/ |
| Mathematica (Version 11) | Wolfram Research | https://www.wolfram.com/mathematica/ |
| Leica Application Suite X | Leica Microsystems | http://www.leica-microsystems.com/products/microscope-software/details/product/leica-las-x-ls/ |
| MicroManager | Vale lab, UCSF | https://micro-manager.org/ |
| GraphPad Prism (Version 5) | GraphPad Software | https://www.graphpad.com/scientific-software/prism/ |
| Adobe Photoshop CS5 | Adobe Systems | http://www.adobe.com/products/photoshop.html |
| Adobe Illustrator CS5 | Adobe Systems | http://www.adobe.com/products/illustrator.html |
| Lasergene Suite (Version 14) | DNASTAR | https://www.dnastar.com/ |

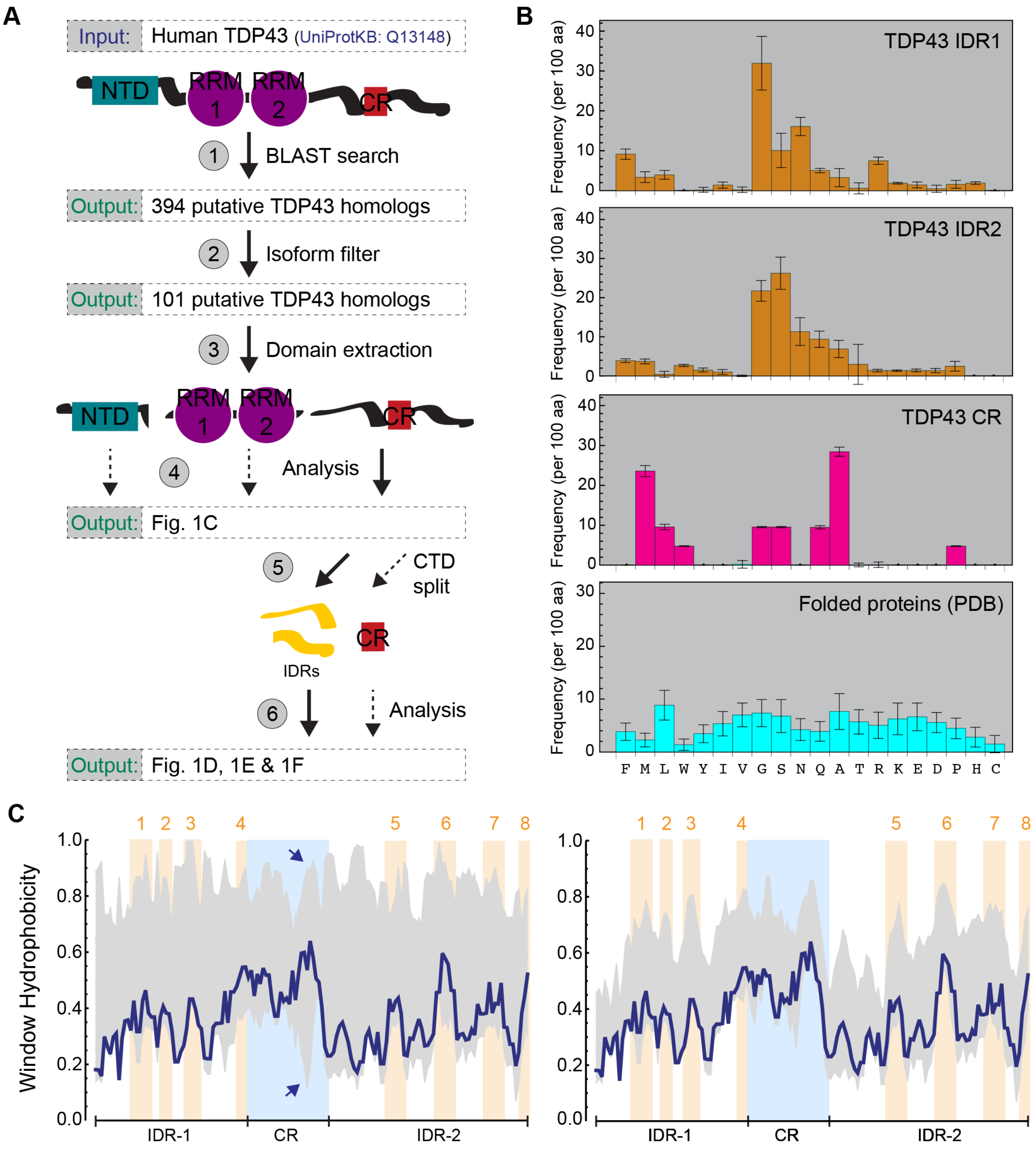


**Supplemental Figure 1 (related to Figure 1).** *Compilation and analysis of a vertebrate TDP43 homolog database*.

(**A**) Human TDP43 (UniProtKB Q13148) was used in a BLAST query to identify vertebrate TDP43 homologs in the non-redundant NCBI protein database (step 1). 394 sequences were initially retrieved and subsequently filtered to retain only a single isoform per species (step 2). The resulting 99 sequences were then divided into separate NTD, RRM and CTD datasets (step 3) for sequence composition analysis (step 4). The CTDs were further split into IDR and CR datasets (step 5) and analyzed as described in Figure 1 and the main text (step 6).

(**B**) Comparison of the amino acid composition of the TDP43 IDRs and CRs to a set of structured proteins from the PDB.

(**C**) Comparison of the CTD hydrophobicity profile using different hydrophobicity scales. Hydrophobicity was calculated in a sliding window of 5 amino acids with the ProtScale tool from ExPASy using different hydrophobicity scales. Blue trace: CTD hydrophobicity according to the scale of Fauchère and Pliska, used throughout the main figures in the manuscript. Yellow shading highlights major hydrophobic clusters. Light gray area: deviation in hydrophobicity profile predictions. The left graph compares scales from: Abraham & Leo, Black, Bull & Breese, Cothia, Eisenberg et al., Guy, Hoop & Woods, Janin, Kyte & Doolittle, Manavalan et al., Miyazawa et al., Rao & Argos, Roseman, Tanford, Welling et al. and Wolfenden et al. There is only one major region of discrepancy (highlighted by the blue arrows), which falls into the CR, which was not altered in this study. The right graph compares predictions from the scales of Abraham & Leo, Black, Cothia, Eisenberg et al., Janin, Kyte & Doolittle, Manavalan et al., Miyazawa et al., Rau & Argos, Roseman and Tanford. Results were re-scaled to range between 0 and 1 for comparison.

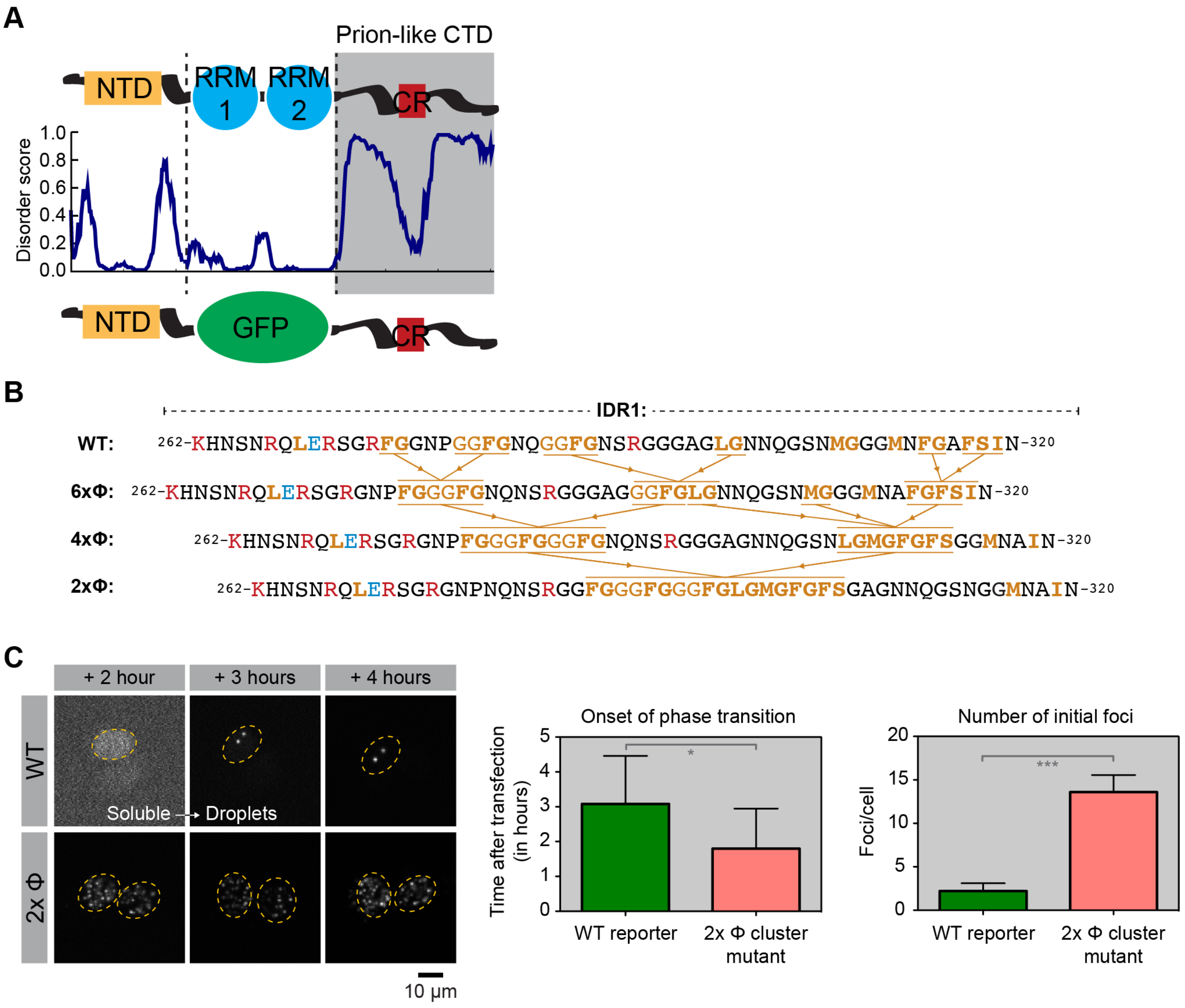


**Supplemental Figure 2 (related to Figures 2, 3 and 4)**. *Design of TDP43$_{RRM\text{-}GFP}$ reporter and CTD mutants with altered spacing between hydrophobic clusters.*

(**A**) Illustration of the TDP43$_{RRM\text{-}GFP}$ reporter design. Note that replacing the RRM domains with GFP leaves the overall domain architecture of TDP43 intact.

(**B**) The spacing between hydrophobic clusters was changed by sliding adjacent sets of clusters together, illustrated for how the WT sequence was changed to the 6xΦ, 4xΦ and 2xΦ variants in IDR1. Hydrophobic motifs are in orange, positively-charged residues in red, and negatively-charged residues in blue.

(**C**) Time course of condensate formation by the WT and 2xΦ variants of TDP43$_{RRM\text{-}GFP}$. Left: Representative images of live cells expressing the WT or 2xΦ variants at the indicated time points after transient transfection. The time of onset for focus formation across multiple cells (center, $n \geq 12$; $p = 0.01$ by Student's t-test) and the number of foci per cell in the first frame after phase separation (right, $n \geq 12$; $p < 0.0001$ in Student's t-test) are shown for the WT and 2xΦ mutant.

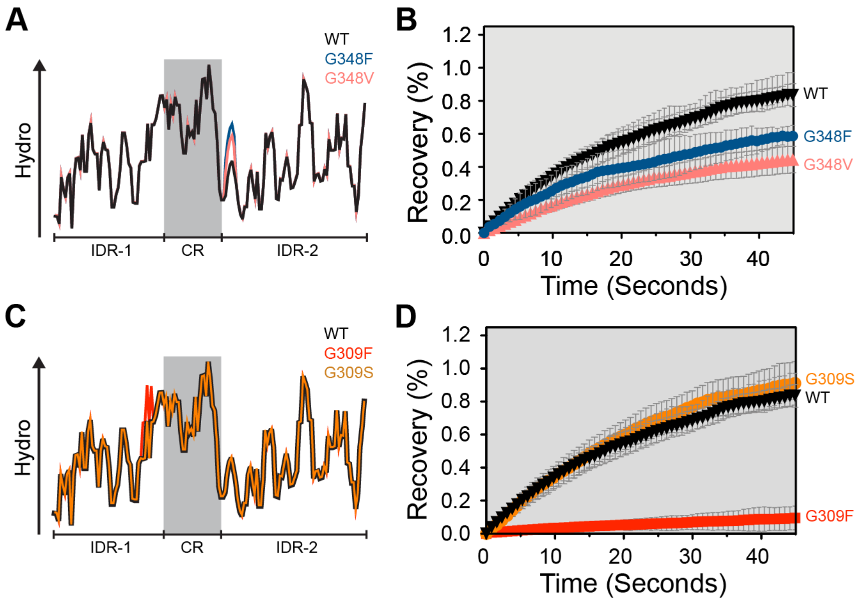


**Supplemental Figure 3 (related to Figure 4).** *Phase behavior of TDP43$_{RRM\text{-}GFP}$ droplets carrying point mutations in IDR1 and IDR2.*

(**A**) The effects of the G348F and G348V mutations on local hydrophobicity predict that both mutations reduce TDP43 phase dynamics.

(**B**) Half-bleach FRAP experiments of G348V and G348F mutant TDP43 droplets show that both mutations reduced fluidity.

(**C**) The effects of the G309F and G309S mutations on local hydrophobicity predict that only the G309F mutation should alter TDP43 phase dynamics.

(**D**) Half-bleach FRAP experiments revealed that the G309F mutation reduced droplet fluidity, while the G309S mutation had no effect.

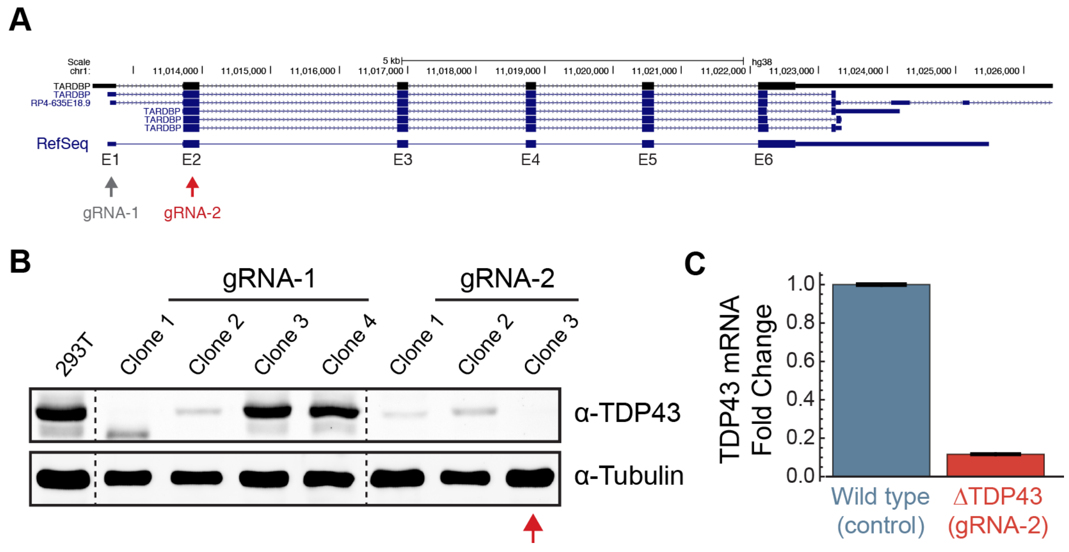


**Supplemental Figure 4 (related to Figures 5, 6 & 7).** *Generation of TDP43-/- HEK-293T cells.*
(**A**) Structure of the *TARDBP* locus (from the UCSC genome browser) showing the exons targeted by two independent sgRNAs.
(**B**) Immunoblot showing TDP43 and α-tubulin (loading control) protein levels in parent HEK-293T cells and clonal derivatives after transient expression of Cas9 and the indicated sgRNAs. The red arrow indicates the clone used to make stable cell lines (clone 3, sgRNA-2).
(**C**) TDP43 mRNA levels were measured using qRT-PCR in HEK-293T cells (WT control) and a clonal TDP43-/- HEK-293T cell line (clone 3, sgRNA-2 in B).

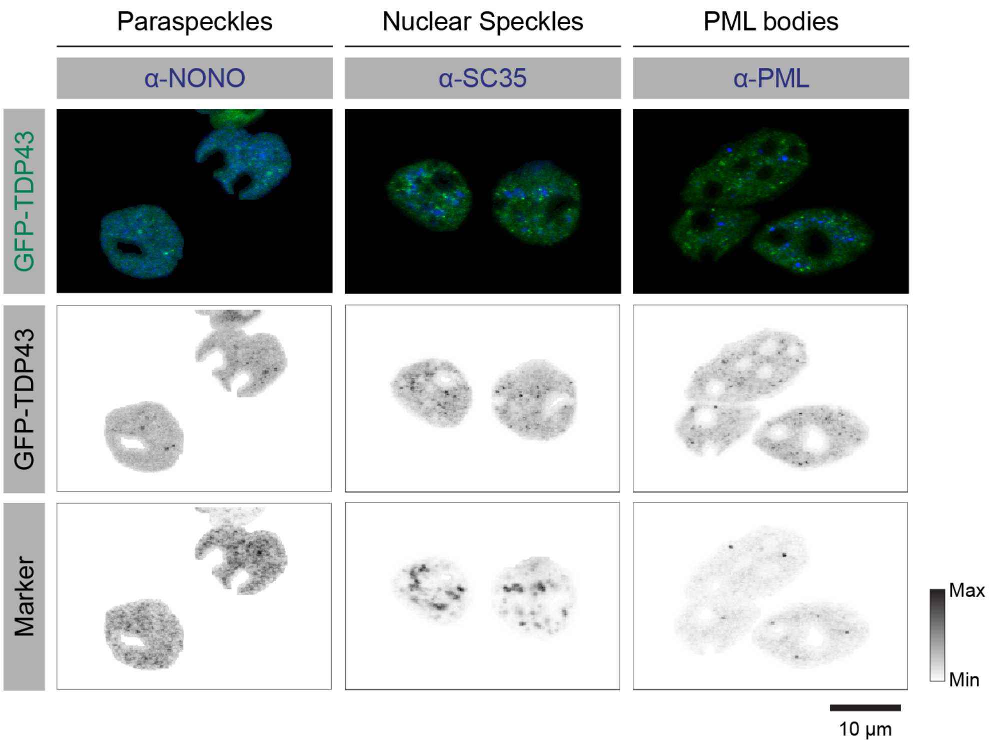


**Supplemental Figure 5 (related to Figure 5).** *GFP-TDP43 foci do not co-localize with paraspeckles, nuclear speckles and PML bodies in HEK-293T cells.*
*TDP43*-/- HEK-293T cells stably expressing GFP-tagged full-length wild-type TDP43 were immunostained using antibodies against the indicated marker proteins to test for co-localization of the nuclear TDP43 foci with known nuclear bodies. ***Top row***: Overlays of representative confocal images showing GFP-TDP43 in green and the indicated marker in blue. ***Middle and bottom rows***: Gray scale heat map representations of the GFP-TDP43 and marker protein signals.

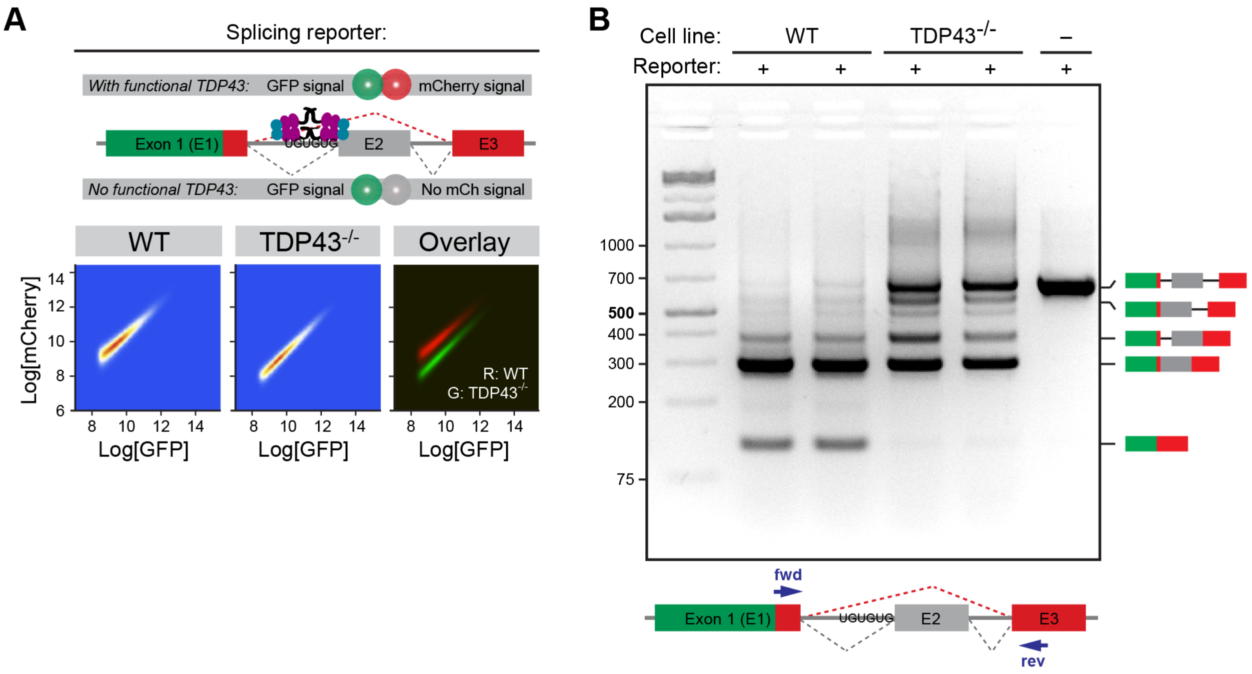


**Supplemental Figure 6 (related to Figure 6).** *Characterization of the TDP43-dependent fluorescent splicing reporter in WT and TDP43-/-* HEK-293T cells*.*

(**A**) Flow cytometry plots showing the distribution of GFP and mCherry fluorescence signals from cells transiently transfected with the splicing reporter shown above. Retention of Exon 2 (E2) will reduce mCherry fluorescence but will leave the GFP signal intact, reducing the mCherry/GFP ratio. Data from WT and *TDP43-/-* cells (left and middle panels) are color-coded as heat maps representing cell density and are overlaid in the right panel for comparison (WT cells in red and *TDP43-/-* cells in green). For each value of GFP fluorescence, the mCherry fluorescence is lower in the *TDP43-/-* cells.

(**B**) RT-PCR analysis to determine the splicing pattern of the reporter in WT and TDP43-/- HEK-293T cells (clone 3, sgRNA2). The cartoon below the gel shows the annealing positions of the primers used in the assay and cartoons to the right of the gel illustrate the structure of various splicing products and intermediates.

```
KHNSNRQLER SGRFGGNPGG FGNQGGFGNS
RGGGAGLGNN QGSNMGGGMN FGAFSINpam
maaaqaalqs swgmmgmlas qqnqsgpSGN
NQNQGNMQRE PNQAFGSGNN SYSGSNSGAA
IGWGSASNAG SGSGFNGGFG SSMDSKSSGW
GM
```

Predicted LARKs
Conserved region

**Supplemental Figure 7 (related to Discussion).** *LARKs are confined to IDR1 of TDP43.* Grey shading indicates the localization of the predicted LARKs in the TDP43 IDRs. The conserved region (lower case letters) is shaded in red. See Discussion for details.
