## Supplemental File 1 for "Decoding and recoding phase behavior of TDP43 reveals that phase separation is not required for splicing function"

**Supplemental File 1.** Comparison of selected tetrapod CTD sequences.

---

Legend:
**Hydrophobic residue**
Conserved region (not mutated)
Region of ambiguity

Aligned CTD sequences:

```
Human      (NCBI Reference Sequence: NP_031401.1)
Mouse      (NCBI Reference Sequence: NP_663531.1)
Xenopus    (NCBI Reference Sequence: AAH61336.1)
Bird       (NCBI Reference Sequence: XP_009865615.1)
```

```
Human      KHNSNRQLERSGRFGGNPGGFGNQGGFGNSR-GGGAGLGNNQGSNMGGG--MNFGAFSINPAMMAAAQAALQSSWGMMGMLASQQNQSG
Mouse      KHNSNRQLERSGRFGGNPGGFGNQGGFGNSR-GGGAGLGNNQGGNMGGG--MNFGAFSINPAMMAAAQAALQSSWGMMGMLASQQNQSG
Xenopus    KHNNNRQLERGGRF-PGPS-FGNQG-YPNSRPSSGA-LGNNQGGNMGGGGGMNFGAFSINPAMMAAAQAALQSSWGMMGMLASQQNQSG
Bird       KHNSNRQLERGGRFGGNPGGFGNQGGFGNSR-GGGGGLGNNQGSNMGGG--MNFGA----PAMMAAAQAALQSSWGMMGMLASQQNQSG

Human      PSGNNQNQGNMQRE-PNQAFGSGNNSYSGSNSGAAIGWGSASNAGSGSGFNGGFGSSMDSKSSGWGM
Mouse      PSGNNQSQGSMQRE-PNQAFGSGNNSYSGSNSGAPLGWGSASNAGSGSGFNGGFGSSMDSKSSGWGM
Xenopus    PQGSNQGQGNQQRDQP-QSFGS-NNSY-GSNSGA-IGWGS-PNAGSGSGFNGGFSSSMESKSSGWGM
Bird       PSGNNQPQGNMQREQ-NQGFSSGNNSYGGSNSGAAIGWGSASNAGSSSGFNGGFGSSMDSKSSGWGM
```
