## Supplemental File 3 for "Decoding and recoding phase behavior of TDP43 reveals that phase separation is not required for splicing function"

**Supplemental File 3.** Sequences of compositional CTD mutants.

---

Legend:
**Hydrophobic residue**
Conserved region (not mutated)

**>WT**
KHNSNRQLERSGRFGGNPGGFGNQGGFGNSRGGGAGLGNNQGSNMGGGMNFGAFSINPAMMAAAQAALQSSWGM
MGMLASQQNQSGPSGNNQNQGNMQREPNQAFGSGNNSYSGSNSGAAIGWGSASNAGSGSGFNGGFGSSMDSKSS
GWGM.

**>Φ-S**
KHNSNRQSERSGRSGGNPGGSGNQGGSGNSRGGGAGSGNNQGSNSGGGSNSGASSSNPAMMAAAQAALQSSWGM
MGMLASQQNQSGPSGNNQNQGNSQREPNQASGSGNNSSSGSNSGAASGSGSASNAGSGSGSNGGSGSSSDSKSS
GSGS.

**>F-S**
KHNSNRQLERSGRSGGNPGGSGNQGGSGNSRGGGAGLGNNQGSNMGGGMNSGASSINPAMMAAAQAALQSSWGM
MGMLASQQNQSGPSGNNQNQGNMQREPNQASGSGNNSYSGSNSGAAIGWGSASNAGSGSGSNGGSGSSMDSKSS
GWGM.

**>FYW-S**
KHNSNRQLERSGRSGGNPGGSGNQGGSGNSRGGGAGLGNNQGSNMGGGMNSGASSINPAMMAAAQAALQSSWGM
MGMLASQQNQSGPSGNNQNQGNMQREPNQASGSGNNSSSGSNSGAAIGSGSASNAGSGSGSNGGSGSSMDSKSS
GSGM.

**>VLIM-S**
KHNSNRQSERSGRFGGNPGGFGNQGGFGNSRGGGAGSGNNQGSNSGGGSNFGAFSSNPAMMAAAQAALQSSWGM
MGMLASQQNQSGPSGNNQNQGNSQREPNQAFGSGNNSYSGSNSGAASGWGSASNAGSGSGFNGGFGSSSDSKSS
GWGS.

**>FYW-L**
KHNSNRQLERSGRLGGNPGGLGNQGGLGNSRGGGAGLGNNQGSNMGGGMNLGALSINPAMMAAAQAALQSSWGM
MGMLASQQNQSGPSGNNQNQGNMQREPNQALGSGNNSLSGSNSGAAIGLGSASNAGSGSGLNGGLGSSMDSKSS
GLGM.

**>VLIM-F**
KHNSNRQFERSGRFGGNPGGFGNQGGFGNSRGGGAGFGNNQGSNFGGGFNFGAFSFNPAMMAAAQAALQSSWGM
MGMLASQQNQSGPSGNNQNQGNFQREPNQAFGSGNNSYSGSNSGAAFGWGSASNAGSGSGFNGGFGSSFDSKSS
GWGF.

**>R-K**
KHNSNKQLEKSGKFGGNPGGFGNQGGFGNSKGGGAGLGNNQGSNMGGGMNFGAFSINPAMMAAAQAALQSSWGM
MGMLASQQNQSGPSGNNQNQGNMQKEPNQAFGSGNNSYSGSNSGAAIGWGSASNAGSGSGFNGGFGSSMDSKSS
GWGM.

**>K-R**
RHNSNRQLERSGRFGGNPGGFGNQGGFGNSRGGGAGLGNNQGSNMGGGMNFGAFSINPAMMAAAQAALQSSWGM
MGMLASQQNQSGPSGNNQNQGNMQREPNQAFGSGNNSYSGSNSGAAIGWGSASNAGSGSGFNGGFGSSMDSRSS
GWGM.

**>KRED-S**
SHNSNSQLSSSGSFGGNPGGFGNQGGFGNSSGGGAGLGNNQGSNMGGGMNFGAFSINPAMMAAAQAALQSSWGM
MGMLASQQNQSGPSGNNQNQGNMQSSPNQAFGSGNNSYSGSNSGAAIGWGSASNAGSGSGFNGGFGSSMSSSSS
GWGM.

**>F-Y**
KHNSNRQLERSGRYGGNPGGYGNQGGYGNSRGGGAGLGNNQGSNMGGGMNYGAYSINPAMMAAAQAALQSSWGM
MGMLASQQNQSGPSGNNQNQGNMQREPNQAYGSGNNSYSGSNSGAAIGWGSASNAGSGSGYNGGYGSSMDSKSS
GWGM.

**>M-S**
KHNSNRQLERSGRFGGNPGGFGNQGGFGNSRGGGAGLGNNQGSNSGGGSNFGAFSINPAMMAAAQAALQSSWGM
MGMLASQQNQSGPSGNNQNQGNSQREPNQAFGSGNNSYSGSNSGAAIGWGSASNAGSGSGFNGGFGSSSDSKSS
GWGS.

**>M-V**
KHNSNRQLERSGRFGGNPGGFGNQGGFGNSRGGGAGLGNNQGSNVGGGVNFGAFSINPAMMAAAQAALQSSWGM
MGMLASQQNQSGPSGNNQNQGNVQREPNQAFGSGNNSYSGSNSGAAIGWGSASNAGSGSGFNGGFGSSVDSKSS
GWGV.

**>6x Φ clusters**
KHNSNRQLERSGRGNPFGGGFGNQNSRGGGAGGGFGLGNNQGSNMGGGMNAFGFSINPAMMAAAQAALQSSWGM
MGMLASQQNQSGPSGNNQNQGNMQREPNQAFGYSSGNNSGSNSGAAIGWGSASNAGSGSNGFGGFGGWGMSSMD
SKSS.

**>4x Φ clusters**
KHNSNRQLERSGRGNPFGGGFGGGFGNQNSRGGGAGNNQGSNLGMGFGFSGGMNAINPAMMAAAQAALQSSWGM
MGMLASQQNQSGPSGNNQNQGNMQREPNQASGNNSFGYSIGWGGSNSGAASASNAGSGSNGFGGFGGWGMSSMD
SKSS.

**>2x Φ clusters**
KHNSNRQLERSGRGNPNQNSRGGFGGGFGGGFGLGMGFGFSGAGNNQGSNGGMNAINPAMMAAAQAALQSSWGM
MGMLASQQNQSGPSGNNQNQGNMQREPNQASGNNSFGYSIGWGGFGGFGGWGMSGSNSGAASASNAGSGSNSMD
SKSS.
